# Notch signaling is a critical initiator of roof plate formation as revealed by the use of RNA profiling of the dorsal neural tube

**DOI:** 10.1101/2020.12.09.417279

**Authors:** Shai Ofek, Sophie Wiszniak, Sarah Kagan, Markus Tondl, Quenten Schwarz, Chaya Kalcheim

## Abstract

The factors underlying establishment of the definitive roof plate (RP) and its segregation from neural crest (NC) and interneurons are unknown. We performed transcriptome analysis at trunk levels of quail embryos comparing the dorsal neural tube at premigratory NC and RP stages. This unraveled molecular heterogeneity between NC and RP stages, and within the RP itself. By implementing these genes, we asked whether Notch signaling is involved in RP development. First, we observed that Notch is active at the RP-interneuron interface. Furthermore, gain and loss of Notch function in quail and mouse embryos, respectively, revealed no effect on early NC behavior. Constitutive Notch activation caused a local downregulation of RP markers with a concomitant development of dI1 interneurons, as well as an ectopic upregulation of RP markers in the interneuron domain. Reciprocally, in mice lacking Notch activity both the RP and dI1 interneurons failed to form and this was associated with expansion of the dI2 population. Collectively, our results offer a new resource for defining specific cell types, and provide evidence that Notch is required to establish the definitive RP, and to determine the choice between RP and interneuron fates, but not the segregation of RP from NC.

**Summary statement:** A new set of genes involved in Notch-dependent roof plate formation is unraveled by transcriptome analysis.

## Introduction

Defining the mechanisms that generate a sequence of specialized cell types and further coordinate their functions is critical for understanding organ development. The dorsal neural tube (NT) epithelium is an excellent system to tackle these questions as it is a highly dynamic structure in both time and space [1]. Initially, it is transiently populated by premigratory neural crest (NC) cells. The latter exit the NT to form the peripheral nervous system as well as a rich assortment of additional derivatives that vary along the neuraxis [2–4]. Subsequently, the dorsal NT midline domain becomes the definitive roof plate (RP) of the spinal cord which is flanked ventrally by dorsal interneuron populations [1,3,5,6].

The RP acts as a neural patterning center that secretes growth factors of the bone morphogenetic protein (BMP) and Wnt families, both crucial for dorsal interneuron development [7–14], and for proliferation of ependymal cells at later stages [15,16].

The morphology of the RP differs along the axis. In the hindbrain, it is composed of a large sheet of cells, whereas more caudally, it is a relatively thin, wedge-shaped strip of cells located along the midline of the spinal cord [17,18]. Fate mapping analysis revealed that the RP generates the choroid plexus at cranial levels of the axis [17,18]. In addition, the RP is transformed into radial glia-like cells [16,19–21], recently found to support growth of spinal cord axons [22]. These radial glial cells eventually generate the stem cell-containing dorsal ependymal zone in the adult spinal cord of humans and rodents [15,16,23].

In spite of its pivotal significance, the origin of the RP in the NT and how it segregates from the earlier NC remained elusive for many years. In fact, the closing dorsal NT was classically termed “RP” yet, essentially, it included the premigratory cohort of NC [1]. To note, several differences exist between these two populations, whereas NC cells actively proliferate, RP progenitors withdraw from the cell cycle [5,24]. Moreover, BMP signaling is needed for NC emigration and for initial RP formation, yet the consolidating RP becomes refractory to BMP activity even if it continues synthesizing the protein [5].

To begin understanding the relationship between NC and RP, we previously implemented spatio-temporally controlled lineage analysis in the avian trunk. This revealed first, that the dorsal NT is sequentially transited by distinct cell populations that emigrate to yield NC derivatives [25,26]; second, that RP progenitors originate ventral to the premigratory NC; and third, that these RP precursors relocate ventro-dorsally to reach their final position in the dorsal midline of the NT upon completion of NC exit [25]. Furthermore, tracing the dynamics of the NC marker *Foxd3* using a specific reporter revealed that NC and RP progenitors are initially part of the *Foxd3* lineage yet RP precursors downregulate *Foxd3* while relocating into the dorsal midline, thus segregating from the NC [26]. The dorsal NT is, therefore, a dynamic area in which progressive NC emigration takes place until replacement by the definitive RP, resulting in the separation between central and peripheral branches of the nervous system [1,25].

The precedent results provided the basis for understanding the transition between NC and RP stages. Nevertheless, a serious limitation for further mechanistic understanding of this process was the lack of differential markers expressed in either population. Whereas several NC specific genes are known (eg; *Foxd3, Sox9*, *Snail2*), many of them (eg; *Lmx1a, Lmx1b, Msx, BMP4, Wnt1*) continue to be expressed in RP, and only few are known to be expressed uniquely in the latter (eg; *cHes1/Hairy1, R-spo1)* [5,27]. For elucidating RP formation, availability of such unique genes is of utmost significance to serve both for identification purposes as well as for functional studies.

In addition to its separation from the NC lineage, a boundary separates the ventral domain of the nascent RP and prospective dorsal interneurons along the spinal cord. Although the RP secretes BMP and Wnt proteins responsible for specification and differentiation of selected interneuron cell types (see above), the factors that determine the establishment of the RP and/or the formation of the boundary between RP and interneurons remain unknown. At cranial levels, maintenance of the boundary between RP and the rhombic lip neuroepithelium was shown to be mediated by Delta (Dll)-Notch signaling but its initial formation was not investigated, neither at cranial levels, nor in the trunk [18]. Notch and its ligands are ubiquitously used for coordination of differentiation between neighboring cells in processes such as boundary formation, lateral inhibition and cell fate determination [28–30].

In the present study, we performed a transcriptome analysis at trunk levels of quail embryos comparing the dorsal NT domain at premigratory NC and RP stages. This analysis yielded genes that were differentially and specifically expressed in either NC or RP. Furthermore, we unraveled a molecular heterogeneity within the trunk-level RP, with transcripts expressed mainly in its lateral or medial domains, respectively. Collectively, our results provide a new resource for investigating a fundamental fate transition during neural development. By implementing the newly described genes in both quail and mouse embryos, we further examined the hypothesis that Notch signaling is involved in *de novo* RP development in the trunk. First, we observed that Notch is active in the dorsal NT except for the dorsal midline region and a similar dorsal limit of mRNA expression was observed for Dll1, Serrate/Jagged, and Lfng, altogether suggesting that Dll-Notch signaling operates at the boundary between the prospective RP and dorsal interneurons. Gain of Notch function in quails revealed a cell autonomous inhibition of RP markers and their replacement by dI1 interneurons. Additionally, an ectopic induction of the expression of RP traits was apparent ventral to the endogenous RP and this was associated with reduced expression of BarHL1, a dI1 interneuron marker. Reciprocally, loss of Notch function was performed in mice by tissue specific knockout of the E3 ubiquitin ligase mindbomb1 (Mib1) to remove the function of all Notch ligands in the dorsal NT and its derivatives. In *Wnt1-Cre; Mib1^fl/fl^* embryos, no RP or dI1 interneurons developed. Concomitantly, dI2-dorsal interneurons expanded dorsalward. In contrast, both gain and loss of Notch function revealed no effect on emigration/migration of NC cells. Together, we provide evidence that Notch signaling is both sufficient and necessary for the choice between RP and interneuron fates without affecting early stages of NC ontogeny.

## Results

### Transcriptional profiling of the dorsal NT at early premigratory NC and RP stages

To conduct a genome-wide transcriptome analysis of dorsal NT development during the transition from NC to RP stages, we electroporated a GFP-encoding plasmid into the dorsal domain of quail NTs at either stage (Supplem. Fig.S1A,B,E,F). Six-to-8 hours later, NTs were isolated, dissociated into single cells, and fluorescent cells were FACS-sorted. No GFP+ cells were apparent in control, non-electroporated samples (Supplem. Fig.S1C,G). In contrast, and as expected from focal transfections, the percentage of live, GFP+ cells was 0.32% and 0.28% of the total input for NC and RP, respectively. In both control and electroporated cases, very low proportions of propidium iodide (PI)+ dead cells were found; these were excluded by gating and only 1000-6000 GFP+/PI− cells/sample were collected (Supplem. Fig.S1D,H).

Next, harvested cells were processed for RNA-sequencing (RNA-seq) in triplicates for each stage (Fig.1A-C). Principal-component analysis (PCA) showed that NC and RP samples cluster into two distinct groups (Fig.1A). Differential gene expression analysis revealed significant up-regulation of 1243 and down-regulation of 1068 genes in RP when compared to NC, respectively, providing a rich data source for further functional analysis (Fig. 1B,C and Supplementary Tables S1 and S2).

**Fig. 1.**
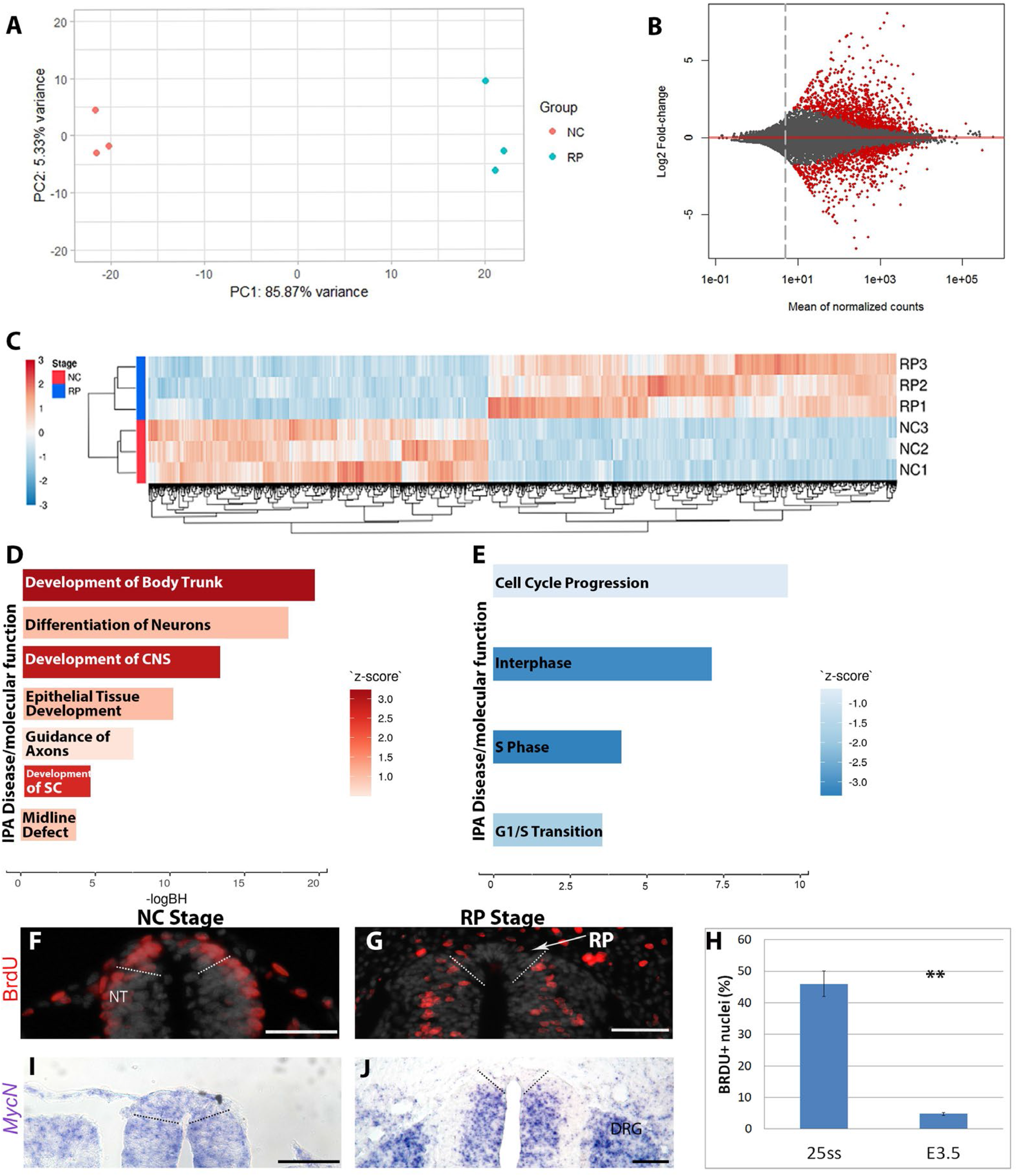
Transcriptome analysis of small groups of premigratory NC and RP cells. (A) Principal component analysis shows a distinct segregation of transcripts for the purified populations of NC and RP cells along 2 principal components axes (PC1, PC2). (B) MA plot comparing RP over NC genes. Genes upregulated in RP when compared to NC are above the red line and downregulated genes are below. (C) Heat map of 2171 differentially expressed genes (padj<0.05) between NC and RP. Red and blue colors represent upregulated and downregulated genes, respectively. Normalized expression is shown after scaling the values for each gene. Genes and samples are ordered by hierarchical clustering, as indicated by the respective dendrograms. (D,E) Ingenuity Pathway Analysis (IPA) showing selected enriched categories that include molecular functions and diseases (p-value<0.05). Categories with IPA Z-scores greater than 2 or smaller than −2 can be considered to be significantly predicted to be up (D, red) or down (E, blue)-regulated, respectively. (F-H) Bromodeoxyuridine (BrdU) incorporation following a 1-h pulse at NC (E2, F) or RP (E3.5, G) stages. Note the presence of the BrdU+ nuclei (red) in NC domain but not in RP (delimited by dashed lines). Quantification of Brdu-positive cells (H) shows significantly reduced number of proliferating cells in the RP (*p<0,001, N=5/stage). (I,J) ISH for MycN shows downregulation in RP (delimited by dashed lines). Abbreviations, DRG, dorsal root ganglion, NT, neural tube. Bar= 50μm.

Further examination by ingenuity pathway analysis (IPA), revealed functional categories containing genes with significant expression changes. Those which were enriched in RP compared to NC (Fig.1D) included “development of the body trunk”, “development of central nervous system” and “development of the spinal cord (SC), reflecting that the RP is a part of the central nervous system. Furthermore, among the RP enriched categories was “epithelial tissue development”, consistent with our previous findings showing that during the transition between NC to RP, the latter regains epithelial characteristics [5]. Enrichment of additional categories such as “guidance of axons” and “midline defects” are also consistent with the established role of the RP as a guidance center for adjacent interneuron projections and that RP abnormalities are a source of NT closure defects, such as spina bifida [31,32]. The presence of “differentiation of neurons” in this analysis may possibly arise because the RP secretes necessary factors for interneuron differentiation [12,33] and/or, due to the inclusion of interneuron progenitors located adjacent to the RP in the sorted cell suspensions.

Ingenuity Pathway Analysis uncovered cell cycle regulation as the major functional group down-regulated in RP compared to NC (Fig.1E). This is consistent with the marked change in proliferative properties, as revealed by BrdU incorporation, between mitotically active premigratory NC compared to the definitive RP, that progressively turns into a post-mitotic structure (45.9±4% vs. 4.8±0.5%, respectively, Fig. 1F-H) [see also [5]]. Along this line, *MycN* mRNA, a marker of proliferative cells [34], was present in the dorsal NT at the NC stage, yet was downregulated in RP (Fig.1I,J). Additional cell cycle associated genes were reduced in RP by at least three-fold, such as *Cdk2-Ap2, olfactomedin4, S100A11, Dlg1, cyclin D3, Rgcc* (Supplementary Table S1); whereas cell cycle inhibitors were up-regulated (*Cdkn2c, Cdkn1c, Dach1* and *2, Btg11, Ndrg1* (Supplementary Table S2).

Taken together, enrichment of the above categories substantiate the validity of our molecular analysis in exposing well-known differential traits of premigratory NC versus RP populations.

### The expression patterns of differential genes

#### NC versus RP markers

To further validate our RNAseq results, we examined by in situ hybridization (ISH) the localization of genes known to be differentially expressed in either NC or RP, as well as that of transcripts never tested before in this context. We present a subset of them alongside graphical representations of their normalized expression based on triplicate samples at either stage. Consistent with our previous results [25,26,35], the NC markers *FoxD3* and *Snail2* were down-regulated at the RP stage (Supplementary Fig. S2A-F). Notably, so were the Wnt inhibitors *Dact2, Sfrp2* and *Dkk1* (Supplementary Fig. S2G-O). In all cases, there was a positive correlation between the transcriptome data and the ISH patterns. Additional transcripts significantly reduced in RP when compared to NC were *Ltk, Mesp1, Runx1, Calponin2, ETS1, ETS2, Glipr2, Ptn, Ripply1, LMO4, Mylk, SALL4, syndecan4, Cdx2, Cdx4-like, Spry2, Nrp2, Tfap2C, Tspan1, Sox18, Mef2C, Gata5* (Supplementary Table S1). Cell adhesion molecules also changed between the stages, with *Cad1, Cad6, Cad11, Cad20, Pcdh1, Pcdh19* and *B-cadherin-like*, being down-regulated in RP. Among up-regulated cell adhesion genes were *Pcdh8, Pcdh9, Pcdh11, Pcdh18, Cad4, Cad8, Cad22, SDK2* (Supplementary Table S2). However, N-Cad (*Cdh2*) mRNA levels did not change, consistent with previous results showing that the dynamics of N-cad function in the dorsal NT is regulated by post-transcriptional processing [36].

Importantly, we found a selection of genes expressed in RP but not premigratory NC (Fig. 2). These included the RP-specific Spondin *Rspo1* [27], and *HES4*, the quail ortholog of chick Hairy1 that exhibits 94% identity at the protein level. In addition, the BMP member *Gdf7*, and the BMP antagonists *BAMBI* and *Gremlin*, the retinoic acid-associated genes *Raldh2* and *CRABP1*, and the chemorepellents of commissural axons *Draxin* and *Slit1* (Fig. 2).

**Fig. 2.**
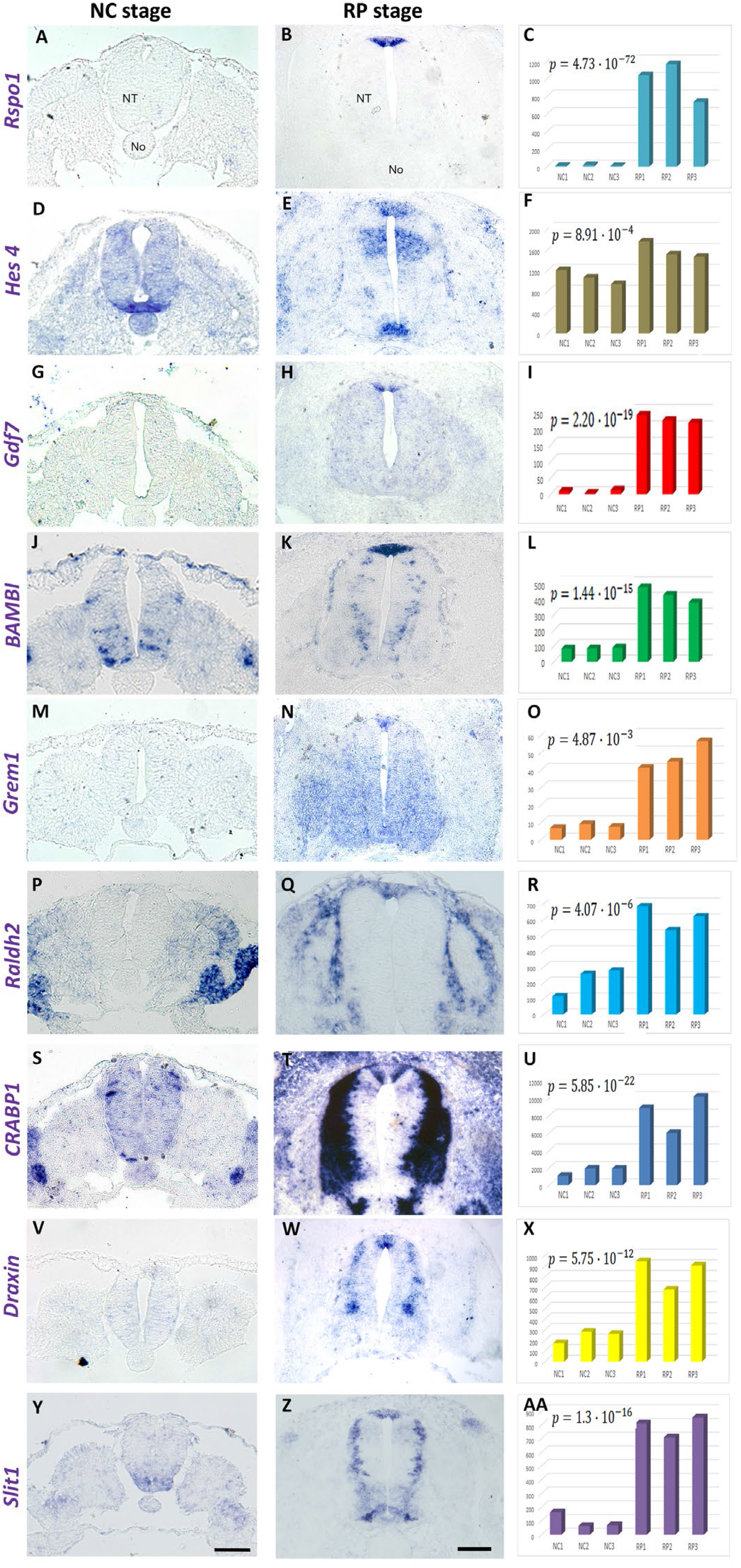
ISH for selected genes upregulated in RP compared to NC. (A, D, G, J, M, P,S,V,Y) ISH at NC and RP (B, E, H, K, N, Q, T,W, Z) stages, respectively. Note expression in the dorsal tube at RP but not NC stage. (C, F, I, L, O, R, U, X, AA) Quantification of gene expression levels stemming from the transcriptome analysis. Note positive correlation between ISH and transcriptome results. In each chart, the B-H adjusted p value is indicated. Abbreviations, NT, neural tube, No, notochord. Bar= 50μm.

In the category of genes enriched in RP compared to NC, there were additional transcripts that exhibited a wider expression pattern that included, but was not restricted to the RP. Examples were *Norrin (NDP), LRP8, Znf536,* and *Zic4* (Supplementary Fig. S3). In all cases, there was also a good correlation between the differential expression patterns and transcriptome results. Together, our RNAseq provided us with a set of spatiotemporal-specific genes appropriate for cell type identification and for functional studies.

#### Expression of “anti RP” markers is complementary to the RP-specific genes

While attempting to validate by ISH the localization of differential transcripts, a subset of genes showing a significant up-regulation in the RP (Supplementary Fig. S4, right panels), yielded a surprising localization pattern. *Dach1, LFNG, CDKN2C, BCL11B, Wnt4, OLFM1* and *INSM1* mRNAs were all expressed in different cell subsets along the NT, except for the RP itself which remained negative (Supplementary Fig. S4). This observation suggests that results of RNAseq cannot be taken as the sole criterion for marker expression in a given cell type, and further spatial confirmation by ISH is warranted.

To define the extent of the nascent RP, we predicted that the precedent genes could prove to be useful if delimiting the ventral boundary of this structure. To this end, we performed ISH of adjacent sections at the RP stage, combining the Notch-related gene *LNFG* with a set of RP-specific transcripts. In all cases examined, the dorsal limit of *LFNG* expression corresponded to the ventral limit of localization of *RSpo1, HES4, BAMBI, Draxin, Raldh2, Gdf7* and *Crabp1*, respectively (Supplementary Fig. S5). Reciprocally, the RP-specific *RSpo1* gene was hybridized in combination with various “anti RP” genes such as *BCL11B, Dach1, OLFM1, INSM1,* and *Wnt4*, exhibiting a comparable complementary pattern (Supplementary Fig. S6, A-O). Hence, the expression of RP and “anti RP” genes enabled us to define the precise size of the RP territory. Measurement of the number of Hoechst-positive nuclei within the RP territory delimited by *LFNG*, revealed that the flank-level RP at E3.5 of quail development consists of 61±1.92 cells per 10μm section (N=4, Supplementary Fig. S6P-R).

#### Molecular heterogeneity within the RP

Careful inspection of the patterns of expression of RP genes, revealed that *Gremlin1, BAMBI, Raldh2, HES4* and *Draxin* were homogeneously distributed throughout the RP (Figs 2 and Supplementary Fig. S5). In contrast, the downstream Wnt target *Axin2,* the Wnt-associated ubiquitin ligase *RNF43*, and the mRNA encoding the RSpo1 receptor *Lgr4* were confined to the medial domain of the RP (Supplementary Fig. S7).

A reciprocal pattern was observed for *RSpo1*, that was primarily transcribed in two peripheral stripes of the RP delimiting the central domains of expression of *Axin2, RNF43* and *Lgr4* (Supplementary Fig. S7) and abutting the ventral limits of the RP (Supplementary Fig. S5A-C and Supplementary Fig. S6). A similar pattern was observed for *Gdf7* [(Supplementary Fig. S5P-R, Fig.2, see also [18]] and *CRABP1* (Supplementary Figs. S5S-U, and Fig.2). Hence, similar to the RP at the hindbrain level [17], the flank-level RP also comprises at least two distinct domains, a medial and two flanking lateral regions (Supplementary Fig. S7J).

### The role of Notch signaling in RP development

Notch activity was found to be involved in maintenance of the hindbrain RP [18]. By implementing a subset of the newly identified genes described above, we investigated whether this signaling pathway plays a role in *de novo* RP formation at spinal cord levels.

#### Expression and activity of Notch ligands and receptors in the dorsal NT

Expression of *Dll1, Jagged1* and *Jagged2* was evident in various domains of the quail NT at E4, except for the RP; thus, localization of ligand mRNAs was complementary to *Rspo1* (Supplementary Fig. S8). *Notch1* was widespread throughout the ventricular layer of the NT, exhibiting an apical localization in RP; and *Notch2* was transcribed in RP (Supplementary Fig. S8).

To examine the activity of Notch, we used an antibody specific to its active intracellular domain (ICD), that recognizes the mouse but not the quail protein. Whereas immunolabeling was almost undetectable at E9.75 (Fig. 3A,B), expression of N1ICD was evident at both E10.5 and E11.5 throughout the NT except for the dorsal-most domain corresponding to the RP (Fig. 3E,F,I,J). To inactivate Notch signaling in the dorsal NT, we conditionally removed the ubiquitin ligase *Mindbomb1 (Mib1*) that is required for the functional internalization of all Notch ligands, by intercrossing *Mib1^fl/fl^* and *Wnt1-Cre* mice [37]. As the Wnt1 promoter drives expression of Cre in the dorsal third of the NT prior to the NC stage, this cross abolishes Notch activity prior to and during RP formation. Indeed, N1ICD immunostaining was specifically absent in the dorsal NT of *Wnt1-Cre; Mib1^fl/fl^*, while still present in more ventral domains (Fig. 3C,D,G,H,K,L), consistent with N1ICD specifically representing Notch activity.

**Fig. 3.**
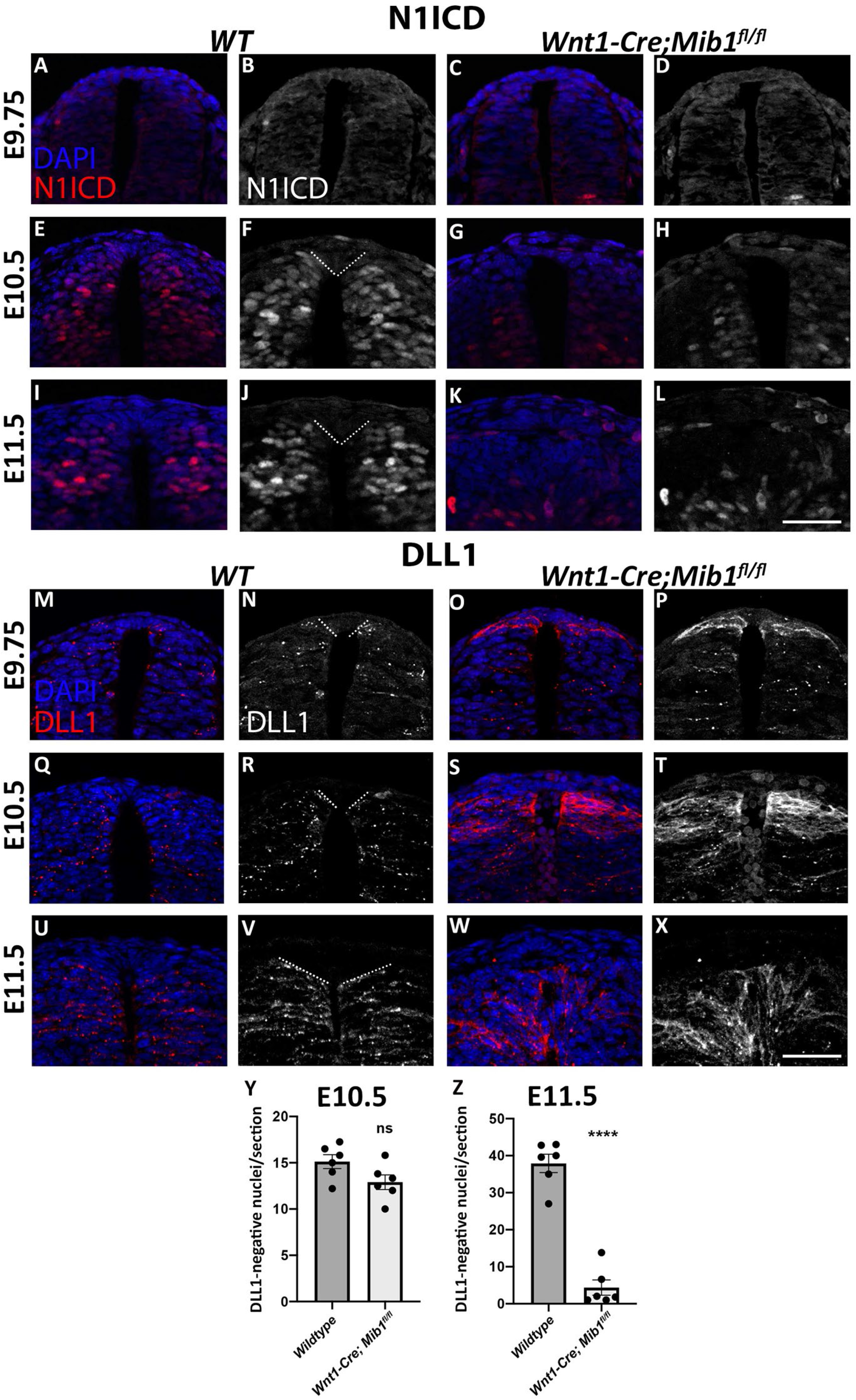
Activation of Notch signaling in the anti-roof plate. (A-L) Transverse sections through the dorsal neural tube at the level of the forelimb in wildtype (WT) and *Wnt1-Cre; Mib1^fl/fl^* mouse embryos, immunostained for N1ICD (white or red) and DAPI (blue). N1ICD staining is shown in combination with DAPI (A, C, E, G, I, K) or as a single color image for clarity (B, D, F, H, J, L). Low levels of N1ICD are evident in the neural tube at E9.75 (A-D), however at E10.5 (E-H) and E11.5 (I-L) N1ICD is broadly expressed throughout the dorsal neural tube in wildtype embryos, whereas this is diminished in *Wnt1-Cre; Mib1^fl/fl^* embryos. (M-X) Transverse sections through the dorsal neural tube at the level of the forelimb in wildtype (WT) and *Wnt1-Cre; Mib1^fl/fl^* mouse embryos, immunostained for DLL1 (red or white) and DAPI (blue). DLL1 staining is shown in combination with DAPI (M, O, Q, S, U, W) or as a single color image for clarity (N, P, R, T, V, X). At all ages examined, in wildtypes, DLL1 is expressed throughout the neural tube except for its dorsal-most region corresponding to the RP (boundary demonstrated by dashed lines in N, R, V). In *Wnt1-Cre; Mib1^fl/fl^* embryos, DLL1 protein accumulates at the cell membrane due to lack of Mib1-induced endocytosis, and by E11.5 there are no DLL1-negative nuclei present in the dorsal-most neural tube. (**Y-Z**) Quantification of DLL1-negative nuclei in the dorsal-most neural tube at E10.5 and E11.5. At E10.5 there is no difference between genotypes, whereas at E11.5 there is a significant reduction in the number of DLL1-negative nuclei, and hence of the presumptive RP, in *Wnt1-Cre; Mib1^fl/fl^* embryos. N=6; ns, not significant; ****p<0.0001. Bar = 50μm

Expression of Dll1 protein exhibited a similar pattern in the wildtype embryos to that observed in quails, with no expression in the dorsal NT and later RP (Fig. 3M,N,Q,R,U,V). As expected, *Wnt1-Cre; Mib1^fl/fl^* embryos revealed enhanced Dll1 signal, consistent with the role of Mib1 in targeting Notch ligands for ubiquitination and their accumulation as inactive ligands in its absence [38]. Most importantly, by E10.5, Dll1 expression shifted dorsally with significantly fewer Dll1-negative nuclei in the dorsal midline region corresponding to the RP, and Dll1 expression reached the dorsal midline by E11.5 with almost no Dll1-negative cells remaining in *Wnt1-Cre; Mib1^fl/fl^* embryos (Fig. 3O,P,S,T,W-Z). Taken together, this indicates that the ventral boundary of the prospective RP, where both ligand and receptors meet, is an area of Notch activity. Furthermore, the dorsalization of Dll1 expression in *Wnt1-Cre; Mib1^fl/fl^* embryos suggests a requirement for Notch signaling in RP formation.

#### Gain of Notch function in quail NTs regulates the choice between RP and dI1 interneuron identities without affecting early NC development

To begin testing the possible involvement of Notch signaling in RP development, we first expressed a constitutively active form of Notch2 (aN2) in embryos aged 25ss, prior to RP formation, followed by fixation at E4 (RP stage). Electroporation of aN2 prevented, in a cell autonomous manner, the upregulation of all the RP markers examined, including *Rspo1, BAMBI, Raldh2,* and *Draxin* (Fig. 4, arrows). In contrast, control GFP had no effect on the expression of either gene (Fig. 4, arrowheads).

**Fig 4.**
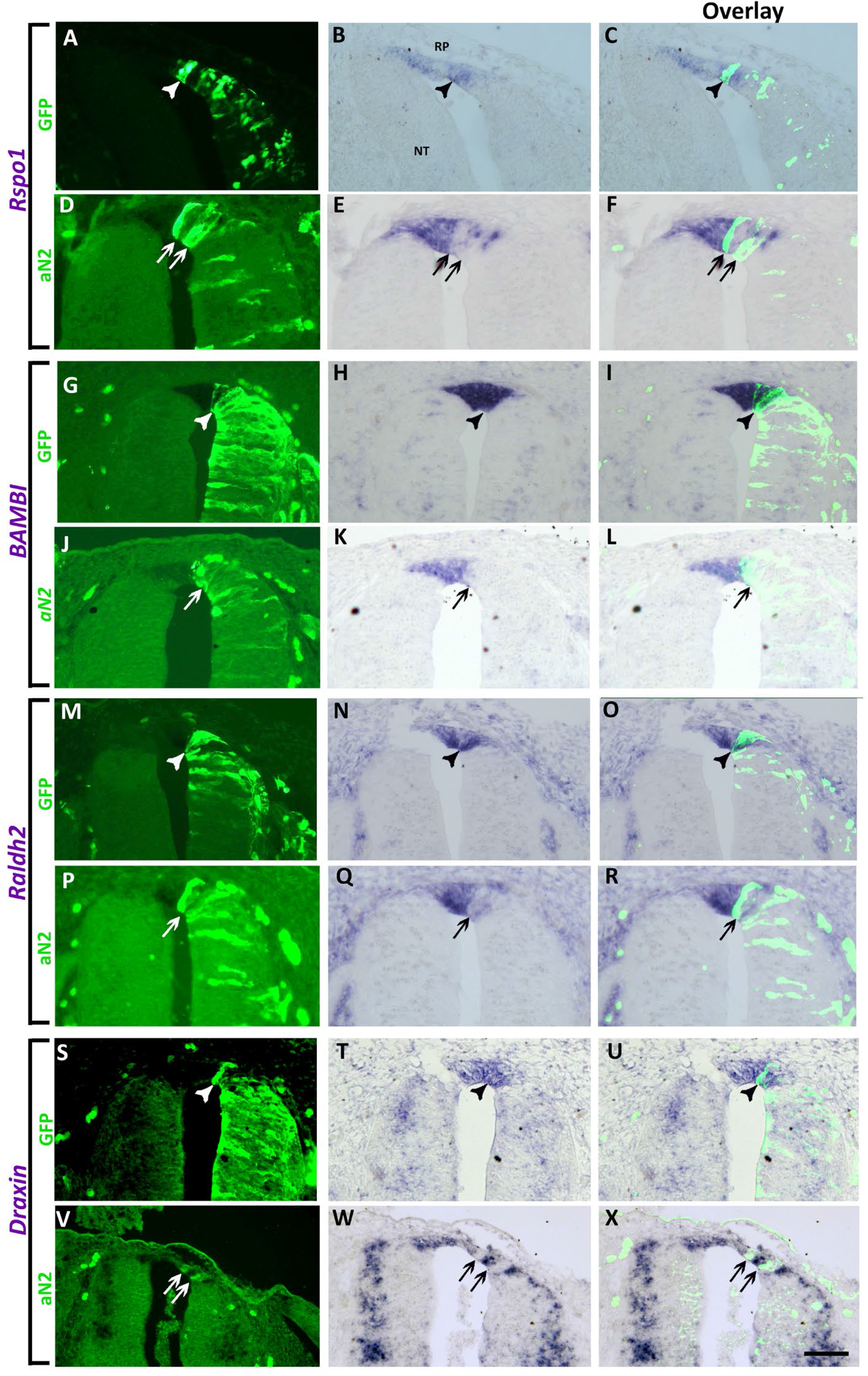
Missexpression of active Notch (aN2) in quail neural tubes prevents the upregulation of RP markers. Left column represents electroporation of control GFP (A,G,M,S) (N=7 for *Bambi,* and N=3 for *Rspo1, Raldh2* and *Draxin*) or aN2-GFP (D,J,P,V) (N=7,15,6,6 for *Rspo1, Bambi, Raldh2* and *Draxin*, respectively.). Middle column depicts ISH for the denoted genes in control GFP (B,H,N,T) or aN2 (E,K,Q,W), and right column is an overlay of the precedent, respectively. Note that control GFP-treated cells co-express both GFP and the RP genes (arrowheads), whereas aN2-treated cells are devoid of marker expression (arrows). Bar= 50μm.

Interestingly, when electroporation of aN2 attained areas adjacent to the RP (Fig. 5, arrows), an ectopic upregulation of *Rspo1, BAMBI, Raldh2* and *Draxin* was often observed more ventrally within the domain populated by dorsal interneurons (Fig. 5, arrowheads). This was not observed upon transfection of control GFP (Fig. 4).

**Fig. 5.**
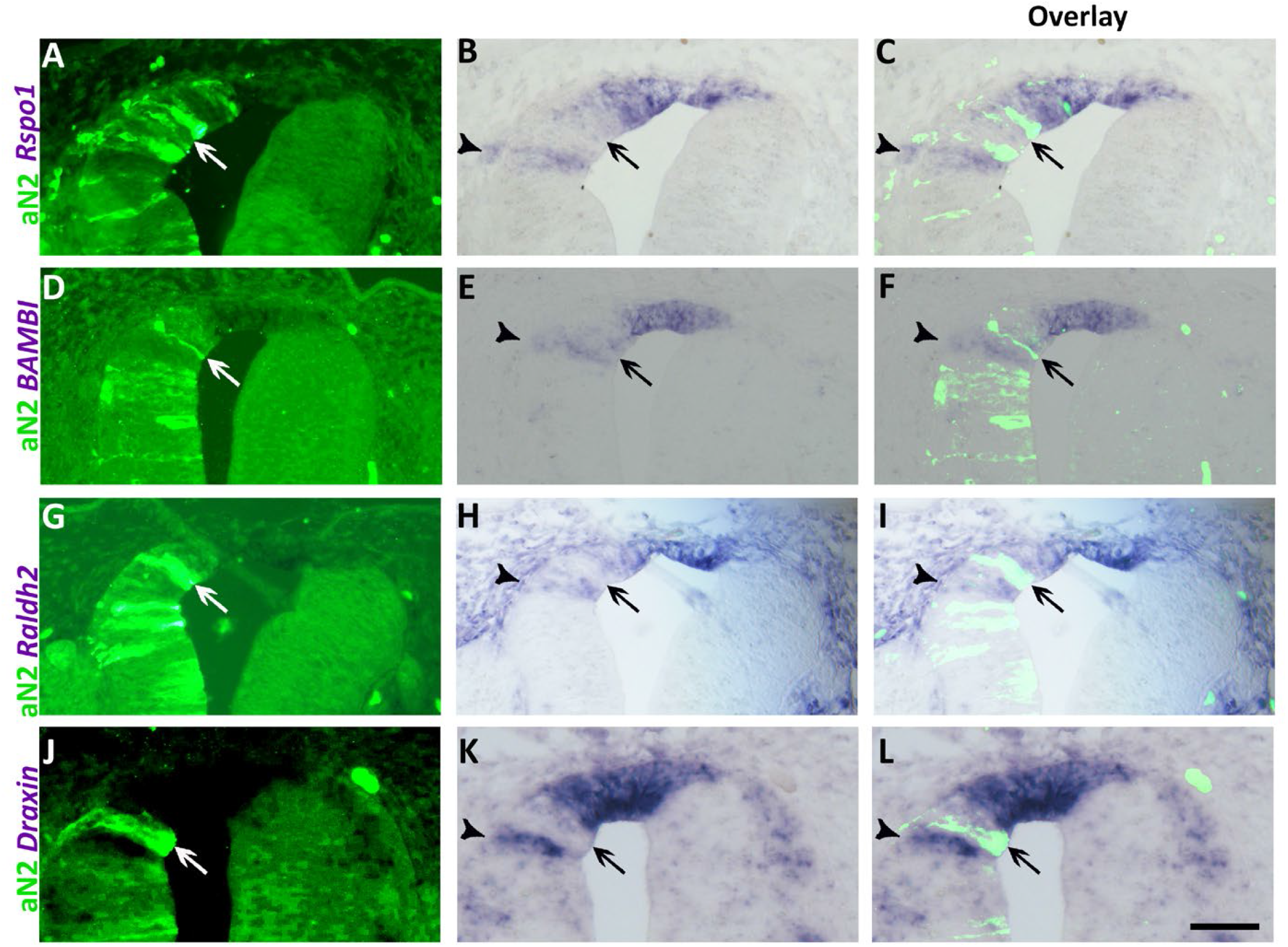
Missexpression of active Notch (aN2) ventral to the prospective RP causes an ectopic upregulation of RP markers. (A,D,G,J) Electroporation of aN2-GFP. (B,E,H,K) ISH of corresponding sections. (C,F,I,L) Overlay of the precedent. Arrows indicate the localization of electroporated cells and arrowheads mark the ectopic upregulation of *Rspo1* (N=6/7), *BAMBI* (N=7/9), *Raldh2* (N=4/6) or *Draxin* (5/5), respectively, within the interneuron domain. For control GFP, see Fig. 4. Bar= 50μm.

These results suggest that ectopic Notch activation in RP suppresses development of RP specific traits while inducing them in the adjacent interneuron domain.

To examine the possibility that interneurons developed at the expense of RP cells, we electroporated control GFP or aN2, and sections were subjected to ISH for *BAMBI* mRNA together with immunolabeling for the dI1 interneuron marker BarHL1. Electroporation of control GFP had no effect on either *BAMBI* expression in RP or on the distribution of BarHL1+ interneurons immediately ventral to the RP (Fig. 6 A-D). In contrast, many dorsal cells missexpressing aN2 lacked *BAMBI* mRNA, and expressed instead the dI1 interneuron marker; as expected, these differentiated neurons localized to the basal domain of the dorsal NT (Fig. 6, E-H, arrowheads). Quantification of BarHL1+ dI1 interneurons within the *BAMBI+* domain confirmed their ectopic development at the expense of RP (Fig. 6 I).

**Fig. 6.**
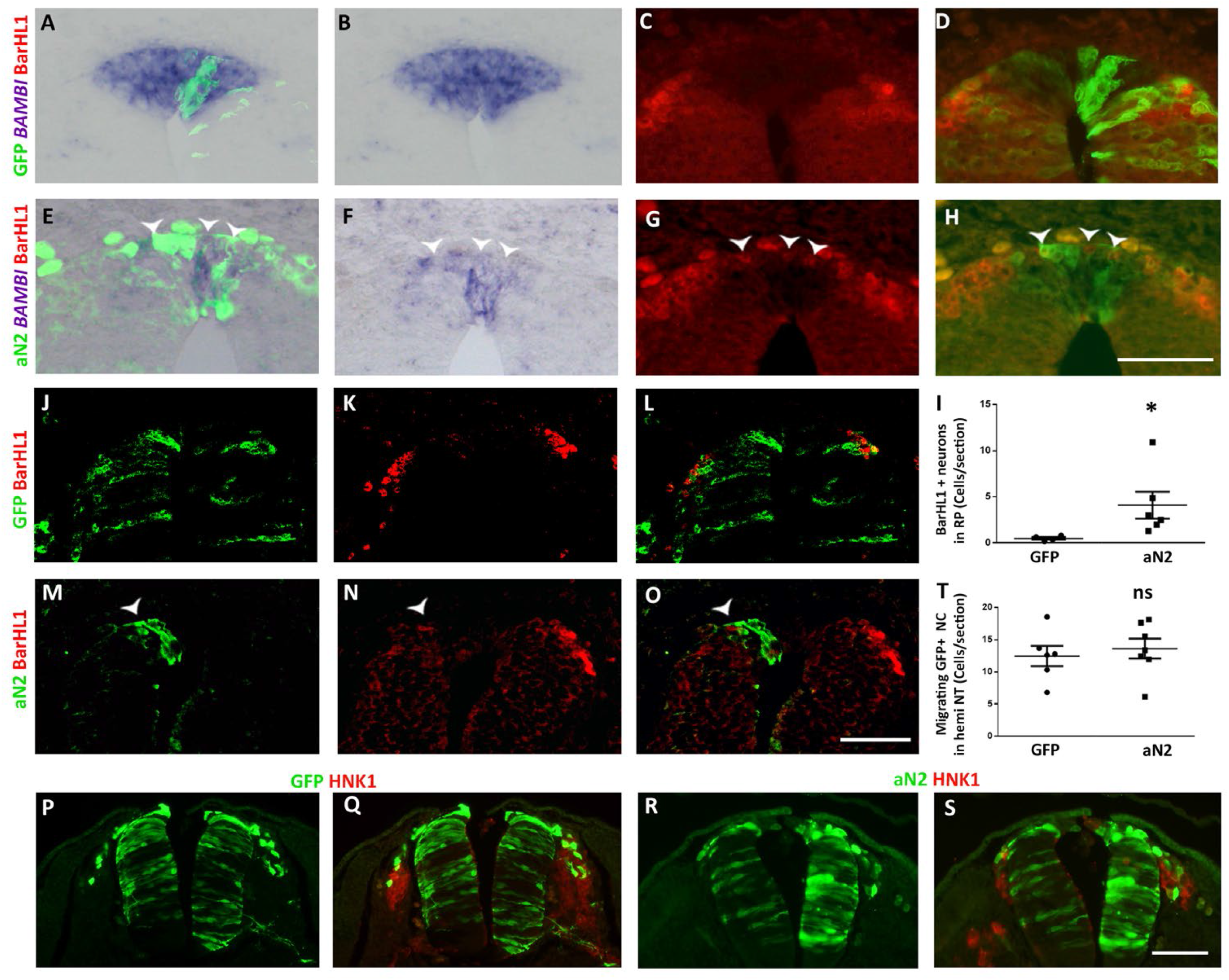
Missexpression of active Notch induces differentiation of dI1 interneurons at the expense of RP traits, yet has no effect on early NC development. (A-D) Electroporation of control GFP. (E-H) Electroporation of aN2-GFP. Note that control GFP has no effect on *BAMBI* mRNA expression (A,B) whereas its upregulation is compromised upon aN2 transfection (E,F). No BarHL1-positive dI1 interneurons are apparent in the RP of control embryos (C,D) (N= 4). In contrast, numerous BarHlL-positive interneurons are present in the RP of aN2-treated cases (G,H, arrowheads). (N= 6). (I) Quantification of the number of BarHL1+ interneurons/section present in the RP, (N=6 for both control and aN2 (*p< 0.02). (J-O) Electroporation of control GFP (J-L) or aN2 (M-O) followed by immunolabeling with BarHL1 antibody. Note that control GFP had no effect on interneuron development (N=4) whereas no BarHL1+ interneurons were apparent in aN2-treated tubes (N=6, arrowhead). (P-S) Gain of Notch activity has no effect on NC migration. Electroporation of control GFP or aN2 at 22ss followed one day later by immunostaining for GFP (green) and the NC marker HNK-1 (red). Note in both cases the migration of GFP+/HNK1+ NC cells. (T) Quantification of the number of GFP-labeled migrating NC cells per section from hemi-neural tubes of control (N=6) and aN2-treated (N=7) embryos (p=0.6, ns). Bar= 50μm.

Next, we examined whether gain of Notch activity outside the RP would affect normal dI1 interneuron development. Control GFP did not alter the expression of BarHL1-positive interneurons, whereas aN2-missexpressing cells lacked BarHL1 immunoreactivity (Fig. 6 J-O, arrowheads), suggesting a failure of dI1 formation upon excess Notch.

At the time of electroporation (22-25ss), NC cells are already specified in the dorsal NT at the trunk level [39]. To further examine a possible effect of aN2 on NC migration, electroporated embryos were fixed a day later. No significant effect was monitored in the number of GFP-positive NC cells in control vs. aN2-treated embryos (Fig. 6P-T). Altogether, misexpression of active Notch affects the choice between adopting a RP fate or a dI1 interneuron identity, yet has no significant effect on emigration or migration of specified NC progenitors (Figs 4-6).

#### Failure of RP formation in mutant mice lacking Notch activity in the dorsal NT

Next, by implementing various RP-specific markers uncovered in the transcriptome analysis of quail embryos, we asked whether Notch signaling is necessary for RP formation in the mouse. Similar to avians, wildtype mouse embryos expressed *Rspo1, GDF7, Raldh2, Slit1*, *Msx1 and Lmx1a* in the trunk RP at E11.5 [(Fig. 7 A,E,G,I,K,M, see also [40]]. At variance with the quail, *BAMBI* was expressed at cranial levels of the mouse axis and was weak to undetectable in the trunk (Fig. 7 C, and not shown). In striking contrast, no expression of the above transcripts was apparent at E11.5 in *Wnt1- Cre; Mib1^fl/fl^* mutants (Fig.7 B,D,F,H,J,L,N). Furthermore, co-labeling of adjacent sections with both *Lmx1a* and Dll1, confirmed the complementary expression of both markers in wildtype embryos, the downregulation of *Lmx1a* in *Wnt1-Cre; Mib1^fl/fl^* embryos and the corresponding dorsalization of Dll1 (Fig. 7M,N). Next, we asked whether the loss of the RP genes was due to a failure in “de novo” RP formation as opposed to its maintenance. To this end, control and mutant embryos were in situ hybridized at E10.5 corresponding to the onset of RP formation, with RP-specific genes not transcribed previously at the NC stage. Expression of *Rspo1, BAMBI, GDF7, Raldh2* and *Slit1* was undetectable in *Wnt1-Cre; Mib1^fl/fl^* mutants when compared to wildtype embryos (Fig. 7 O-X). Together, our data demonstrate that Notch activity in the dorsal NT is necessary for initiation of RP formation.

**Fig. 7.**
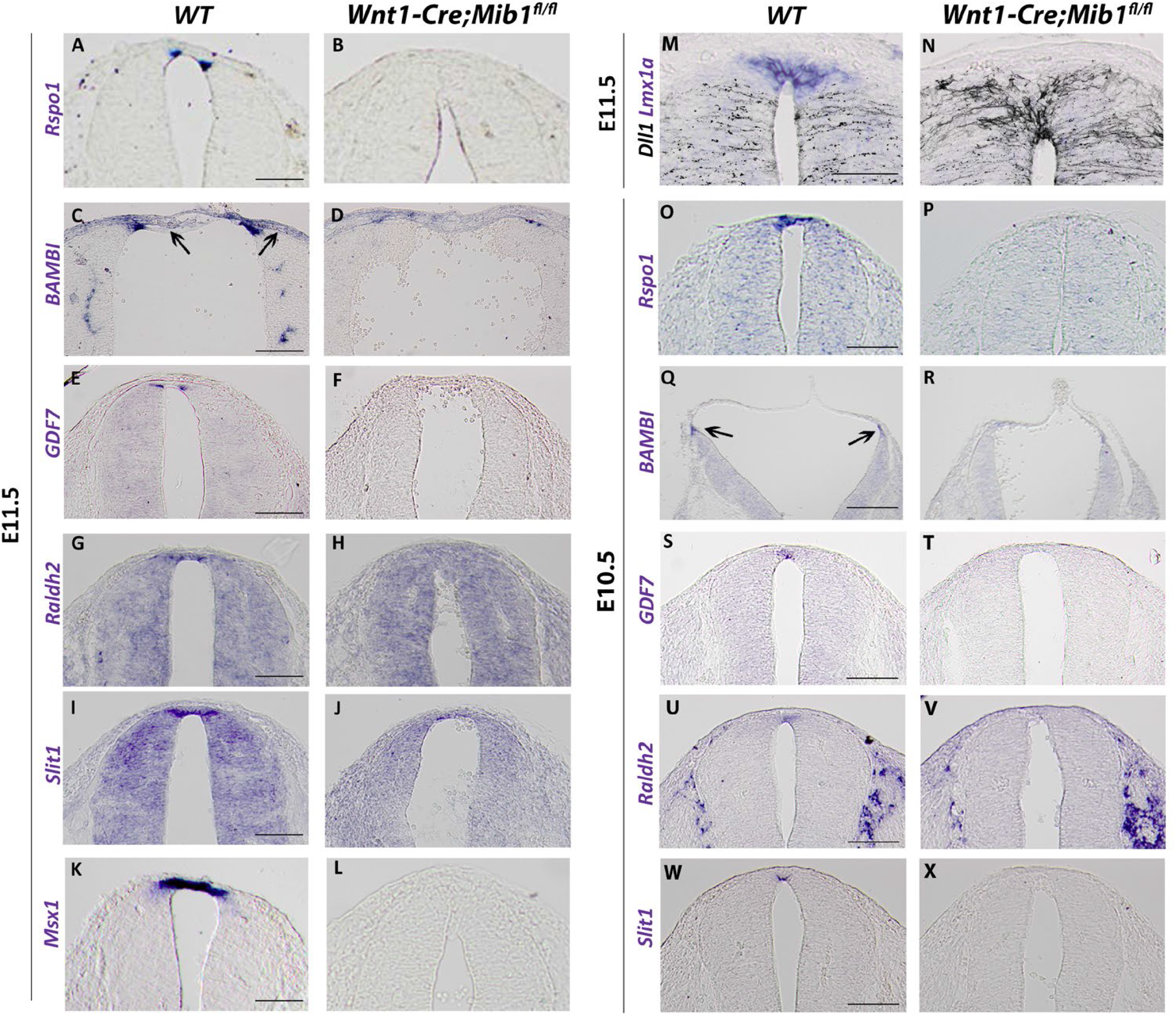
Lack of roof plate formation in the absence of Notch signaling. *In situ* hybridization of definitive RP markers on transverse sections through the dorsal neural tube at the level of the forelimb (A,B, E-N, O,P, S-X) or hindbrain (C, D, Q, R) in wildtype (WT) and *Wnt1-Cre; Mib1^fl/fl^* mouse embryos at E11.5 and E10.5. Expression of all RP *in situ* hybridization markers are missing from mutants at E11.5 and E10.5: *Rspo1* (A, B and O, P); *BAMBI* (C, D and Q, R); *GDF7* (E, F and S, T); *Raldh2* (G, H and U, V); *Slit1* (I, J and W, X); *Msx1* (K, L). (M, N) Combination of adjacent sections of *Lmx1a in situ* hybridization (blue) and immunostaining for Dll1 (black) showing complementary expression of the two markers at E11.5 in WT embryos but loss of *Lmx1a* in mutants with a corresponding dorsalization of Dll1. Bar= 100μm.

#### Loss of Notch results in absence of RP and dI1 interneurons

It has been previously shown, that in the absence of a RP, development of dI1, the dorsal-most interneurons, is abnormal [41,42]. We examined whether Notch-dependent failure of RP formation is associated with a similar phenotype. *Math1*-positive dI1 interneuron progenitors were located lateral to the RP in E10.5 and E11.5 embryos. In contrast, in *Wnt1-Cre; Mib1^fl/fl^* embryos, very few *Math1+* progenitors remained at E10.5 and none were detected at later ages (Fig. 8A-D). Hence, in the absence of a RP, the formation of dI1 interneurons is compromised.

**Fig. 8.**
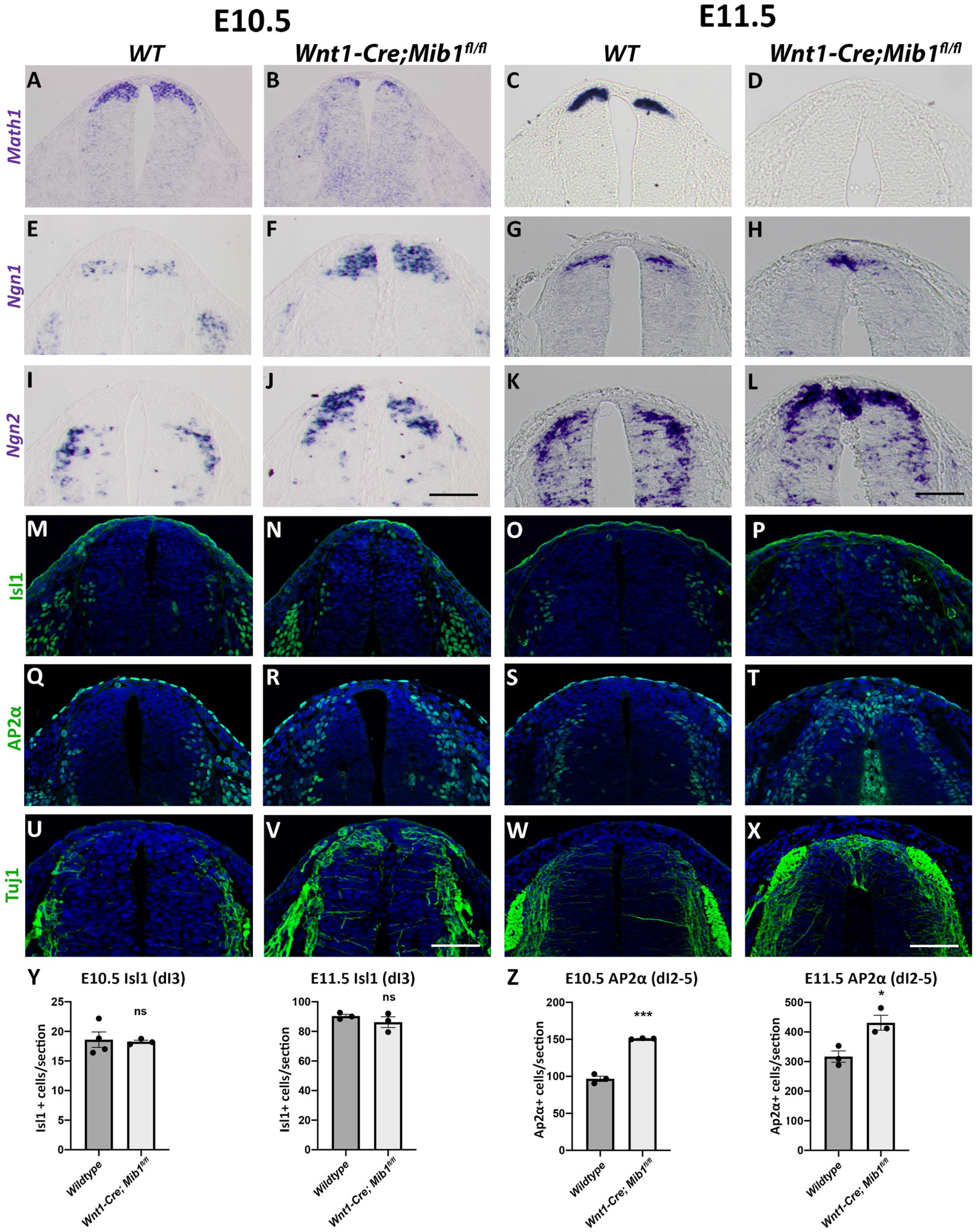
Re-specification of the dorsal neural tube domain in the absence of Notch signaling. (A-L) *In situ* hybridization of transverse sections through the neural tube at the level of the forelimb in wildtype (WT) and *Wnt1-Cre; Mib1^fl/fl^* mouse embryos. *Math1* expression at E10.5 (A, B) and E11.5 (C, D) showing loss of dI1 population in mutants; *Ngn1* expression at E10.5 (E, F) and E11.5 (G, H) and *Ngn2* expression at E10.5 (I, J) and E11.5 (K, L) showing dorsal-ward expansion of the dI2 population. (M-X) Immunostaining of transverse sections through the dorsal neural tube at the level of the forelimb in wildtype (WT) and *Wnt1-Cre; Mib1^fl/fl^* mouse embryos. Isl1 expression at E10.5 (M, N) and E11.5 (O, P) showing a dorsal-ward expansion of dI3 interneurons at E11.5; AP2*α*expression at E10.5 (Q, R) and E11.5 (S, T) showing a dorsal-ward expansion in mutants at both stages (positive staining within the neural tube lumen in T is due to erythrocyte autofluorescence); Tuj1 expression at E10.5 (U, V) and E11.5 (W, X) showing ectopic expression of the axonal marker Tuj1 in the dorsal neural tube of mutants. (Y-Z) Quantification of dI interneuron subtypes in the dorsal neural tube at E10.5 and E11.5. Isl1-positive dI3 interneuron numbers are unchanged in *Wnt1-Cre; Mib1^fl/fl^* embryos (Y), whereas AP2*α*-positive dI2-5 interneuron numbers are increased in *Wnt1-Cre; Mib1^fl/fl^* embryos at both E10.5 and E11.5 (Z). N=3-4 embryos; ns, not significant; *p=0.022; ***p=0.0001. Bar = 50μm.

Next, we monitored expression of *Ngn1+* and *Ngn2+* dI2 interneuron progenitors and observed that their localization in the mutants progressively shifted towards the dorsal midline at the expense of the missing dI1 population. Although ISH of these markers prevented us from accurate quantification, a clear increase in the dI2 interneuron subset was apparent at both stages (Fig. 8 E-L), suggesting aberrant specification of dorsal progenitors into dI2 neurons. Notably, Isl1+ dI3 interneurons were also shifted dorsal-ward in *Wnt1-Cre; Mib1^fl/fl^* embryos, but their total amount was unaffected (Fig. 8 M-P,Y). Further visualization of AP2*α−*positive dI2-5 interneurons [40], revealed a similar dorsal shift at both stages, with their total amount being significantly enhanced in *Wnt1-Cre; Mib1^fl/fl^* embryos at E10.5 and E11.5, respectively (Fig. 8Q-T,Z). Together, this increase is likely to be accounted for by expansion of the dI2 population at the expense of dI1 interneurons, since the Islet1+ dI3 interneurons were not quantitatively affected. Consistent with the replacement of RP by interneurons, we further documented a remarkable dorsalization of the neuronal marker Tuj1 in *Wnt1-Cre; Mib1^fl/fl^* embryos, that was already apparent at E10.5 and progressively reached the dorsal midline by E11.5 (Fig. 8, U-X).

As also previously reported, we observed no effect on specification and emigration of AP2*α*+ NC cells at E9.5 in *Wnt1-Cre; Mib1^fl/fl^* mutants when compared to wildtype [(Supplementary Fig. S9 A,B, and see [37] for quantification], suggesting that Notch signaling is involved in RP/interneuron development but not in early stages of NC ontogeny, as also seen in our quail experiments.

In addition, we asked whether the loss of RP and dI1 interneurons was due to enhanced cell death. No evidence for cleaved caspase3-positive cells was found in the dorsal NT of *Wnt1-Cre; Mib1^fl/fl^* embryos at E9.5, E10.5 or E11.5 when compared to control embryos. In contrast, NC-derived dorsal root ganglia revealed significant cell death at E11.5, as previously documented (Supplementary Fig. S9 A-F) [37]. Likewise, we tested for cell proliferation of the Dll1-negative dorsal cell population by pulse labeling E10.5 wildtype and *Wnt1-Cre; Mib1^fl/fl^* embryos with EdU. About 40% of prospective RP progenitors still proliferated at this stage in wildtype embryos. A 30% reduction in the proportion of EdU+ nuclei was measured in the mutants compared to controls (Supplementary Fig. S9 G-I). Reduced proliferation in the dorsal NT of *Wnt1-Cre; Mib1^fl/fl^* embryos at E10.5 coincides with the replacement of RP progenitors with post-mitotic interneurons (Figs. 7,8).

Altogether, in the absence of Notch signaling, initial NC development was normal, yet no RP or dI1 interneurons developed. This loss could not be explained by changes in cell death or proliferation; instead, it is likely to be accounted for by aberrant specification of the dorsal NT with corresponding expansion of the dI2 interneuron population.

## Discussion

Understanding the role/s of gene activity in the transition between cellular states and fate decisions is essential for elucidating mechanisms of normal development as well as the origin of embryonic malformations. The dorsal NT provides an excellent platform for investigating the sequential formation and delamination of a variety of NC cell types followed by the end of NC production/emigration and the concomitant establishment of the definitive RP [1]. Although the formation and emigration of the NC has been extensively studied [2,3,43], how the dorsal NT moves from the early NC all the way into the definitive RP stage remains largely unknown. This is mainly due to the scarcity of RP-specific markers that enable tracking of the shift in cell identity and understanding the molecular pathways involved in the process. Using RNA-seq and ISH, we identify genes differentially expressed at either stage in the trunk level of the axis, their detailed expression patterns, and infer below on their possible functions in the above transition.

The observed coverage patterns correlate well with the known functions of BMP signaling in the proliferation, delamination and emigration of NC cells [39,44] and, in particular, with the reported end of BMP activity that leads to the arrest of NC emigration and RP formation [5]. The latter process is illustrated by the up-regulation of the BMP inhibitors *BAMBI, Grem1, Norrin, Smad3, PRDM12*, and also that of *HES4*, the quail ortholog of Hairy1/HES1, shown to inhibit BMP signaling in RP [5]. Furthermore, synthesis of factors of the BMP family shown to be necessary for adjacent interneuron development, such as Gdf7 [10], or BMP4 and BMP7 [9,13] is either up-regulated or maintained in RP, respectively. Hence, we further extend the molecular repertoire in support of the notion that the RP becomes refractory to BMP signaling whereas interneurons continue responding to this family of RP-derived factors.

Interestingly, the observed up-regulation of the BMP inhibitor *BAMBI*, raises the possibility that the retinoic acid (RA) pathway is also involved in the end of NC production and/or in RP formation. First, BAMBI is a direct target of RA [45], second, the RA biosynthetic enzyme *Raldh2* begins to be transcribed in RP in addition to its earlier synthesis in paraxial mesoderm (Fig.2) and so are *CRABP1* that binds and mediates RA activities (Fig.2) [46] and the RA modulator *Znf536* [47]. Together, we posit that RA emanating from the RP may contribute to the end of NC production by inhibiting BMP signaling, at least partially through BAMBI, a hypothesis that will be put to the test.

One of the major genes produced selectively in RP but not premigratory NC is *R-spo1*. All four secreted R-spondins are unique agonists of Wnt-mediated signaling [48] which plays a fundamental role in G1/S transition of NC cells and their delamination [44], yet whose possible function in RP formation was so far not explored. R-spondins bind to Lgr 4/5/6 receptors and bring these into position to interact with the ubiquitin ligases Rnf43/Znrf3. This interaction results in membrane clearance of the latter and consequent persistence of the Wnt/Frizzled/Lrp complexes in the plasma membrane, enhancing Wnt signal strength and duration [49,50]. R-spo1 was reported to be involved in male sex reversal and adult intestinal stem cell homeostasis [48] yet its role in RP development remains unknown. Since its expression in RP was compromised in *Wnt1/3a* double knockout mice, it was suggested that it may contribute to the development of dorsal NT under the regulation of Wnts [27]. Consistent with such a possibility, our RNA-seq unravels the up-regulation of the R-spo1 receptor *Lgr5*, induction of *Norrin*, a R-spo/Wnt agonist that additionally inhibits BMP activity [51], and a decrease of *Rnf43*. Moreover, several Wnt inhibitors are down-regulated in RP, such as *Dkk1, Frzb, DACT2* and *Sfrp2*. In contrast, the Wnt receptors *Fz3* and *Fz10*, the downstream target *Axin2* as well as the co-receptors *LRP 4,8,* and *11*, are enhanced.

Our data show that *R-spo1, Gdf7* and *CRBP1* are predominantly expressed in the lateral RP along the trunk [our data and see [18,27]] abutting the boundary between the RP and adjacent neuroepithelium. In contrast, *Axin2, Lgr4* and *Rnf43*, all belonging to the R-spo/Wnt module (see above), are expressed in the medial RP, suggesting that the trunk RP is molecularly heterogeneous. Indeed, this molecular heterogeneity also seems to be conserved in mice where *R-spo1* also marks the lateral RP domains. Apart from uncovering molecular components of this heterogeneity in the trunk, we confirm the notion suggested for the hindbrain RP, that the latter is formed by molecularly and temporally separable lineages [[17,18], and refs. therein]. Importantly, at cranial levels, only the lateral RP seems to contribute to the formation of the choroid plexus [17]. Whether the observed heterogeneity in the trunk RP predicts a differential fate and/or function of medial vs. lateral domains remains to be elucidated.

Taken together, results of our transcriptome data enabled us first, to postulate new roles and interactions between candidate signaling pathways and specific markers of RP domains in the dynamics of dorsal NT development, and second, to plan concrete experiments to put these to the test.

Along this line, we presently addressed the role of the highly conserved Notch signaling pathway in RP formation. In both avians and mice, the dorsal limit of Notch ligand expression corresponds to the ventral boundary of the RP. Although *Notch* mRNAs are widely transcribed, analysis of N1ICD reveals a similar dorsal limit of activity as that of its ligands. This suggests that the ventral boundary of the prospective RP is a region of Notch/Dll signaling.

Mosaic electroporation of active Notch in prospective RP prevented the advent of RP-specific markers in a cell autonomous manner, while inducing them in ectopic positions immediately ventral to the RP in a non-cell autonomous manner. Reciprocally, loss of endogenous Notch activity prevented *de novo* RP formation. Hence, both gain and loss of function approaches unravel a role for Notch signaling in development of the definitive RP.

This interpretation was possible by implementing RP-specific markers, not expressed in the earlier premigratory NC population. Previous studies used BMP4, Wnt1 and Lmx1 as RP markers [52] yet these are transcribed in both populations.

Of importance is the finding that gain of Notch function in the dorsal NT, prevented the expression of endogenous RP traits, while upregulating ectopically specific markers of dI1 interneurons. Conversely, loss of RP development in the absence of Notch signaling compromised development of *Math1*+ dI1 interneurons, a phenotype that was coincident with expansion of the dI2 population. Likely, this may be associated with our observation that Notch activity in normal mouse embryos corresponds to the dorsal limit of dI1 interneurons. These results show: first, that Notch plays a role in the binary choice between RP and interneuron fates, and second, confirm that the RP is necessary for at least dI1 interneuron development [41,42]. Alternatively, changes in Notch activity in the boundary between RP and interneurons may affect both populations altogether, as previously suggested for the hindbrain where a well-defined boundary zone may act as a bidirectional signaling center [18].

It is noteworthy that different experimental paradigms affecting the advent of a definitive RP have varied outcomes to dorsal interneurons. For instance, ablation of the RP by targeting diphtheria toxin under the regulation of *Gdf7* caused the disappearance of both dI1 and dI2 interneurons and the dorsal midline was occupied instead by dI3 cells [42]. In *Lmx1a* mutants, however, *Math1+* dI1 interneurons were generated albeit in reduced numbers and no effect on dI2 was monitored [41]. This difference in phenotype severity could be accounted for by the documented lack of Gdf7, BMP6, Msx1 and Wnt1 in the Gdf7 mutants compared with the Dreher mice in which residual expression of Wnt1 and Msx1/2, factors shown to operate on interneuron development persisted [52,53].

In our loss of Notch function experiments, dI1 progenitors, as assessed by *Math1* expression, were reduced albeit still apparent at E10.5, and absent by E11.5; dI2, but not dI3 interneurons, were expanded in their absence. It is possible that early specification of interneuron subsets occurs prior to the advent of the definitive RP by dorsal NT-derived BMPs [9,12,54]. Additionally, it is possible that BMPs derived later from the ectoderm dorsal to the spinal cord [55], still enable development of selected interneurons of the dI2 subclass, expanded in our mutants. In this context, it would be interesting to examine the relationship between Notch signaling and dorsal NT factors of the BMP and Wnt families.

In both avians and mice, the protocols implemented did not affect early development of NC cells. In avians, epidermal Dll1 signaling was shown to induce NC cells indirectly via BMP activation [56]. In the present study, however, electroporation was performed following initial NC specification and no overt defects in subsequent migration were monitored. Nevertheless, at later stages, Notch was shown to play significant roles in neuron-glia fate decisions within NC-derived dorsal root ganglia [37,57].

Taken together, the combination of RP-specific markers with targeted gene misexpression allows us to begin understanding the mechanisms underlying segregation of the three sequential lineages that inhabit the dorsal NT, e.g; NC, RP and interneurons. Of the precedent lineages, we report that Notch signaling is required for RP formation and for the choice between RP and interneuron fates, yet is dispensable for early stages of NC ontogeny.

## Acknowledgments

We thank Ilana Blech for technical assistance with the avian part of the study; Tamar Hashimshony, Tal Katz-Ezov and Olga Karinsky from the Genome Center at the Technion, for assistance with the RNA seq. We are indebted to Sharona Elgavish, Yuval Nevo and Hadar Benyamini for help with bioinformatic analysis, and Avihu Klar for helpful suggestions. We also thank Xiangjun Xu for technical assistance with mouse genotyping.

## Author Contributions

CK and QS conceived the project and all authors designed the experiments; SO performed the transcriptome analysis and all experiments in quail embryos; SW performed the mouse embryos experiments; SK and MT assisted with probe preparation and ISH analysis; and CK wrote the manuscript. All authors discussed and agreed on the results and approved the manuscript.

## Data availability

Raw RNAseq files will be deposited to GEO after acceptance for publication.

## Materials and Methods

### Quail and mouse embryos

Fertilized quail (Coturnix coturnix Japonica) eggs were obtained from commercial sources (Moshav Mata) and incubated at 38℃ to the desired developmental stages. All experiments on mice were approved by the SA Pathology and University of South Australia Animal Ethics Committee (project # u26/20). To obtain embryos of defined gestational ages, mice were mated in the evening, and the morning of vaginal plug formation was counted as embryonic day (E) 0.5. Pregnant dams were humanely euthanized at relevant days post vaginal plug detection by CO_2_ inhalation and cervical dislocation. To delete *Mib1* specifically in the dorsal neural tube, we mated *Mib1^fl/+^* males carrying a heterozygous *Wnt1-Cre* transgene (Danielian et al., 1998) to *Mib1^fl/fl^* female mice [59]. At least 3 embryos/age/genotype were analyzed for each experiment.

### Expression vectors and *in ovo* electroporation

For transcriptome analysis, pCAGGS-EGFP (2–5 µg/µl) was microinjected into the lumen of the NT opposite the segmental plate region of embryos aged 18ss to label the early premigratory NC or at the flank region of embryos aged 43ss to label the RP. An inducible version of the intracellular part of Notch2 (aN2) [60] was sub-cloned into a tetracycline-sensitive plasmid (pBI-TRE-GFP) [61] as previously described [62]. pCAGGs and pBI-TRE-GFP were used as controls. To perform focal electroporations directed to the dorsal NT at each stage, a 5 mm tungsten electrode was inserted under the blastoderm (negative electrode) and a fine 2 mm long electrode was placed dorsal to the neuroepithelium. For hemi-NT transfections, electrodes were placed at both sides of the NT. A square wave electroporator (BTX, San Diego, CA, USA) was used to deliver one to three pulses of current at 10–20 V for 8 ms.

### Cell dissociation

Dorsally electroporated NT’s were dissected manually 6-8 hours after electroporation before significant emigration of NC cells begun in the early stage embryos. Prior to NT isolation, embryos were inspected under a fluorescent binocular. Embryos with emigrating cells, or with large electroporated domains were excluded. Single cell suspensions were obtained by enzymatic digestion with 0.01% Trypsin/0.02% EDTA, 7µg/ml DNAse1 in Ca/Mg-free PBS for 3.5 and 4.5 minutes for NC and RP stages, respectively. Dissociation was completed by manual pipetting. The resulting cell suspensions were centrifuged at 1000 RPM in an Eppendorf centrifuge, washed once with washing solution (Ca/Mg free PBS/10%FCS/0.1 units of RNAse inhibitor), and then with fluorescent activated cell sorting (FACS) buffer (PBS CaMg free /2% FCS). The final suspension was filtered through a 40µm pore size Cell strainer.

### FACS sorting

ARIA III (BD Biosciences) was used for sorting. The cut-off for sorting the electroporated GFP+ cells and for exclusion of dead/damaged PI+ cells was chosen manually. Three samples for each stage containing a total of 1000-6000 GFP+/ propidium iodide (PI)-negative cells were collected in Trizol. As expected from very focal transfections, the percentage of GFP+/PI− cells of the total population was about 0.3% at both stages. Total viability of the cells was 99.9% and at least 95% or 87% for the electroporated GFP+ cells at NC and RP stages, respectively.

### Library preparation

RNA was extracted according to standard Trizol protocol (Invitrogen), using LPA as carrier (Sigma). Amount and quality were checked on Agilent Bioanalyzer using an RNA-pico chip. 120 pg RNA of each sample was taken for library preparation using the CEL-Seq2 protocol [63], with the following modifications: RNA was added to dNTP/ERCC spike-in mix, and 3 different CEL-Seq primers added to each sample as technical replicates. Library was sequenced on HiSeq2500 in rapid mode, 15 bases for read 1 and 50 bases for read 2.

### Bioinformatic analysis

Demultiplexing was performed according to the pipeline from CEL-Seq2 [63], R2 reads were split into their original samples using the CEL-Seq barcode from R1. The reads were cleaned using trim_galore (uses cutadapt version 1.10) for removal of adaptors, polyA, low-quality sequences (Phred<20) and short reads (<25bp, after trimming). Mapping to the Coturnix coturnix Japonica genome was conducted using Tophat2 version 2.1.0. The reference and annotation files were downloaded from: https://www.ncbi.nlm.nih.gov/genome/term=Coturnix+japonica, ftp://ftp.ncbi.nlm.nih.gov/genomes/refseq/vertebrate_other/Coturnix_japonica/latest_assembly_versions/GCF_001577835.1_Coturnix_japonica_20, respectively. HTseq-count version 0.6.1 was used for counting. Normalization and differential expression analysis using DESeq2 R package version 1.14.1 was performed after combining reads of the technical replicates.

### Proliferation assays and Immunohistochemistry

#### Quail embryos

Incorporation of BrdU was performed 1 hour before fixation and immunolabeling was done as described [5]. For immunostaining, embryos were fixed in 4% formaldehyde overnight at 4°C, processed for paraffin wax embedding and sectioned at 8μM. Antibodies used were rabbit anti BarHL1 (1:100, Sigma, HPA004809), rabbit anti GFP (1:1000, Invitrogen, Thermo-Fisher Scientific, A6455) and rabbit anti RFP (1:000, OriGene, AP09229PU-N). Nuclei were visualized with Hoechst.

#### Mouse embryos

To examine cell proliferation, EdU (5-ethynyl-2’-deoxyuridine) (Life technologies) was prepared in 0.9% NaCl solution at 10 mg/ml. EdU was delivered by intraperitoneal injections to pregnant dams at E10.5 at 100 mg/kg of body weight and embryos were collected 1 hr after injection.

Embryos were fixed in 4% formaldehyde overnight at 4°C, cryopreserved in 20% sucrose, and embedded in OCT compound for cryosectioning. Sections were blocked in 10% Dako serum-free blocking reagent in PBS /0.1% TritonX-100, followed by incubation in primary antibody for 2hr at room temperature or overnight at 4°C. Fluorescent Alexafluor-conjugated secondary antibodies were incubated for 1h at room temperature. Sections were mounted in Prolong Diamond antifade with DAPI. The following primary antibodies were used: mouse anti-Tuj1, 1:750 (Sigma-Aldrich T5076); mouse anti-Isl1, 1:50 (DSHB 40.3A4); rabbit anti-cleaved Caspase-3, 1:500 (Cell Signaling Technology 9661); mouse anti-AP2*α*, 1:20 (DSHB 3B5); sheep anti-Dll1, 1:200 (R&D Systems AF5026); rabbit anti-N1ICD, 1:100 (Cell Signaling Technology 4147). Sections of EdU-labeled embryos were incubated with Click-iT EdU Kit Alexa-555 conjugated (Life Technologies) following staining with primary antibodies.

### Antigen retrieval

Immunostaining for interneuron markers and N1ICD required antigen retrieval. Paraffin sectioned embryos were immersed in pre-boiled sodium citrate pH6, 0.05% Tween20, and microwaved for 10 min (20% strength). Blocking was performed with 5% newborn calf serum (Biological Industries 04-102-1A, Israel) in PBS for one hour at room temperature. Cryosections were incubated in 10mM sodium citrate pH6.0 at 90°C for 20min, and then cooled to room temperature before proceeding with immunohistochemistry procedure as above.

### Tyramide amplification

Immunostaining for N1ICD required tyramide signal amplification, performed using Invitrogen Alexa Fluor 555 Tyramide SuperBoost Kit (B40923). Following primary antibody incubation, sections were incubated with goat anti-rabbit horseradish peroxidase, then tyramide signal amplification was performed for 30 minutes, following manufacturer’s instructions.

### In situ hybridization

*Quail embryos*-ISH was performed as described previously [36]. Embryos were fixed in Fornoy (60% ethanol, 30% formaldehyde, 10% acetic acid), then dehydrated in ethanol/toluene, processed for paraffin wax embedding and sectioned at 10 *μ* m. Slides were sequentially rehydrated in toluene/ethanol/PBS, treated with proteinase K (1µg/ml, Sigma Aldrich P2308) at 37°C for 7 minutes, and then fixed in 4% formaldehyde at room temperature for 20 minutes. Next, slides were washed in PBS followed by 2X SSC and hybridized in hybridization buffer (1X salt solution composed of 2M NaCl, 0.12M Tris, 0.04M NaH_2_PO_4_2H2O, 0.05M Na_2_HPO_4_, 0.05M EDTA, pH7.5), 50% formamide, 10% dextran sulfate, 1mg/ml Yeast RNA, 1X Denhardt solution) containing 1μg/ml DIG labeled RNA probes (prepared with a DIG RNA labeling mix, Roche, 11277073910) for overnight at 65°C in a humid chamber. Post-hybridization, slides were rinsed in a rotating incubator with 50% formamide, 1X SSC, 0.1% Tween 20, until coverslips dropped and then an additional wash for 1 hour followed by 2 washes in MABT (10% Maleic acid 1M pH 7.5, 3% NaCl 5M, 0.1% Tween 20) and preincubation in MABT/ 2.5% FCS. Anti-DIG-AP antibody (1/1000, Roche 11093274910) diluted in MABT+2% BBR+20% FCS was then added for overnight at room temperature. This was followed by rinsing in MABT and then in NTMT (2% NaCl 5M, 10% Tris HCl 1M pH9.5, 5% MgCl2 1M, 0.1% Tween20), and then incubation in NTMT + 1:200 NBT/BCIP Stock Solution (Sigma-Aldrich, 11681451001) at 37°C until the AP reaction was completed.

The following probes were kindly provided: *Foxd3* [64,65], *Snail2* [66], *Dact2* [67], *Sfrp2, Dkk1, LFNG,* and *Wnt4* [68]. Additional probes were produced by PCR amplification (using Q5 high fidelity DNA polymerase) with specific primers (see Supplementary. Table S3 for list of primers). Template for PCR was cDNA, which was synthesized by RNA precipitation followed by reverse transcription PCR. RNAs were produced from 20ss-E4 quail embryos. Tissue samples were homogenized with TriFast reagent, and RNA was separated with chloroform and isopropanol.

#### Mouse embryos

Section ISH was performed as described [69]. Riboprobes were transcribed from plasmids containing PCR amplified cDNA sequences (Supplementary Table S3). m*Ngn1*, m*Ngn2, and Msx1* riboprobes were kindly provided by F. Guillemot. These and *Math1* were prepared as previously described [37,70].

### Data analysis and statistics

The number of Isl1+ or AP2*α*+ nuclei in the dorsal NT was monitored in at least 5 sections per wildtype or *Wnt1-Cre; Mib1^fl/fl^* embryos. The number of BarHl1-positive interneurons or of migrating GFP-positive NC cells was measured in at least 10 sections per embryo from control GFP or aN2-electroporated embryos. Graphs represent mean number of positive cells/section ± SEM.

For measurement of BrdU incorporation, cells in 15 sections per embryo were counted and expressed as percentage of BrdU+/total GFP+ cells in the dorsal NT.

For monitoring cell proliferation in mice, the number of Edu-positive cells was counted in nuclei located dorsal to the boundary of Dll1 expression, ie. Dll1-negative nuclei, and expressed as a percentage of total Dll1-negative nuclei. At least 5 sections per embryo were quantified. Graphs represent mean number of positive nuclei ±SEM. Images were photographed using a DP70 (Olympus) cooled CCD digital camera mounted on a BX51 microscope (Olympus). For figure preparation, images were exported into Photoshop CS6 (Adobe). If necessary, the levels of brightness and contrast were adjusted to the entire image and images were cropped without color correction adjustments or γ adjustments. Data were analyzed using unpaired Student’s *t* test. A *p*-value ≤ 0.05 was considered significant.

## Legends to Supplementary Figures and Tables

**Supplementary Fig. S1.**
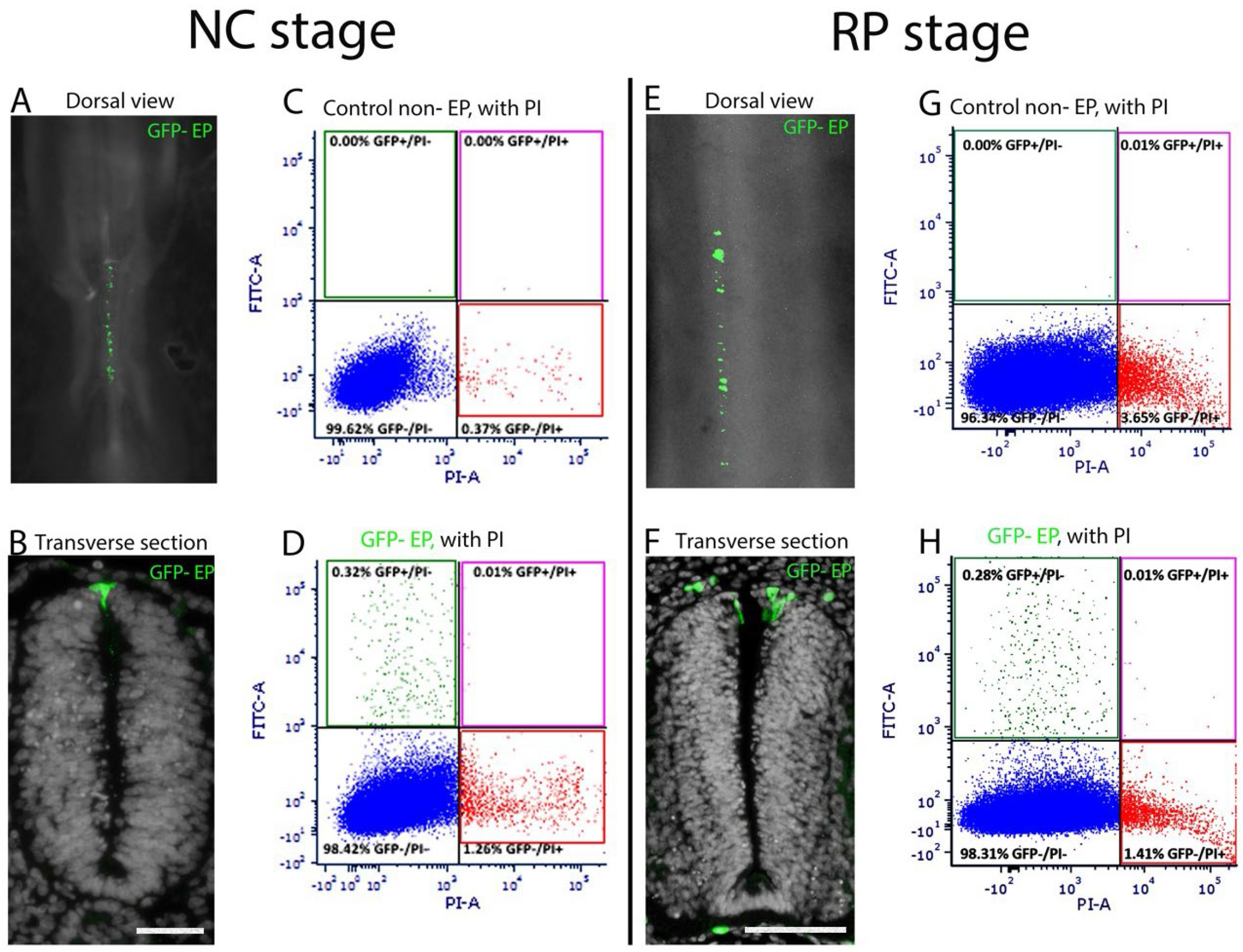
Labeling of the dorsal NT at NC and RP stages followed by FACS analysis. Focal electroporations of GFP-DNA were directed to the dorsal NT at NC and RP stages. Embryos were sacrificed 6-8hr later to isolate premigratory NC progenitors prior to the onset of EMT. (A,E) dorsal views of live embryos showing restricted labeling of the dorsal midline domain at both stages. (B,F) Focal dorsal labelings confirmed in transverse sections. Green cells outside the NT in F represent autofluorescence of blood cells. (C,D,G,H) FACS purification of GFP-labeled cells. Note absence of GFP+ cells in control non-electroporated samples (C and G) and very low proportions of propidium iodide (PI)+ dead cells in both control and electroporated cases. PI+ cells were excluded by gating and GFP+/PI− cells were collected (D,H). As expected from focal transfections, the percentage of live, labeled cells (GFP+/PI−) was 0.32% and 0.28% of the total input for NC and RP, respectively. Abbreviations, EP, electroporation, FITC, fluorescein isothiocyanide. Bar= 50μm.

**Supplementary Fig. S2.**
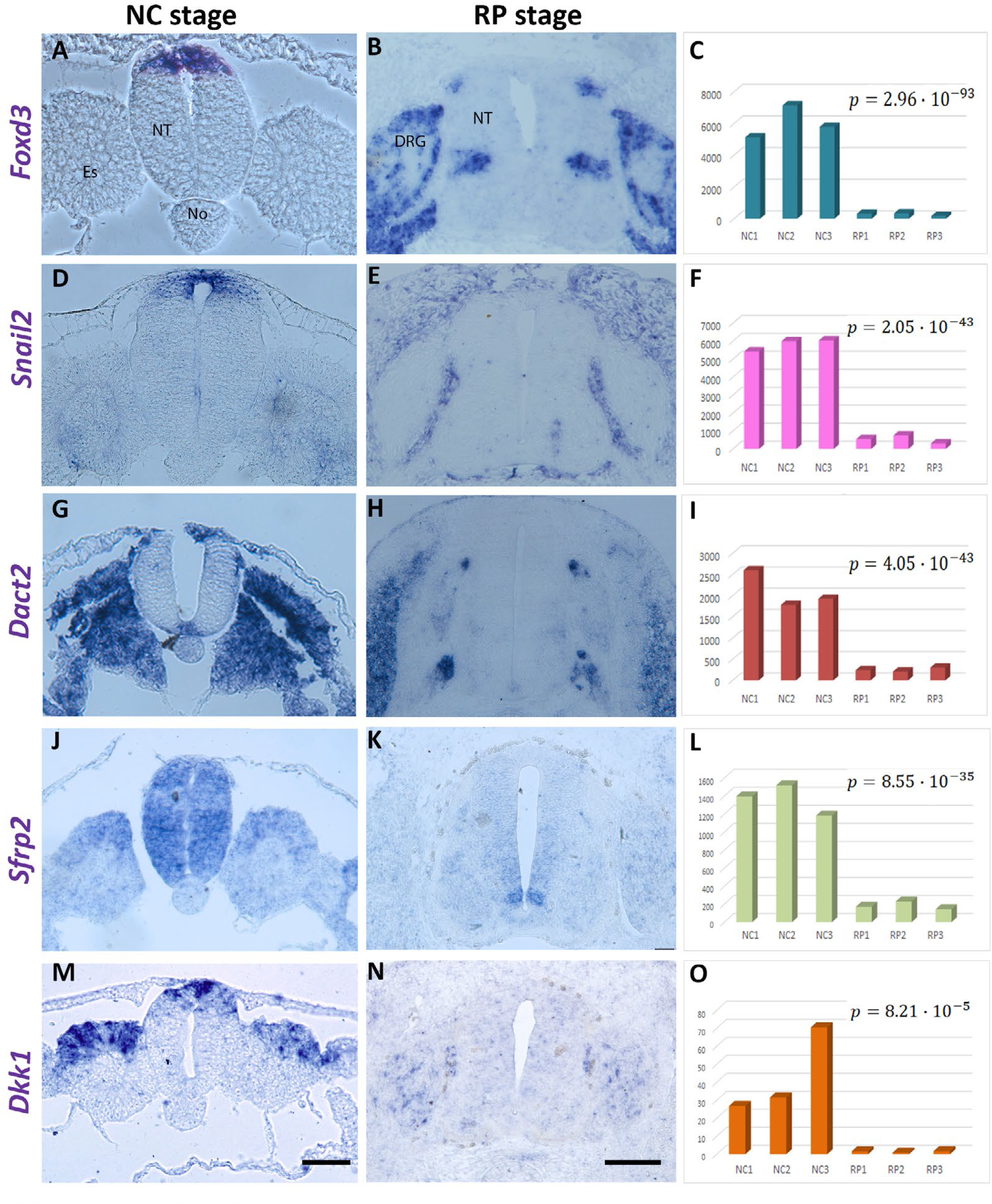
In situ hybridization (ISH) for selected genes downregulated in RP compared to NC. (A, D, G, J, M) Transcripts expressed in the dorsal tube at the NC but not RP stage (B, E, H, K, N). (C, F, I, L, O) Quantification of gene expression levels stemming from the transcriptome analysis. NC1-3 and RP1-3 represent experimental triplicates. Note positive correlation between ISH and transcriptome results. In each chart, the B-H adjusted p value is indicated. Abbreviations, DRG, dorsal root ganglion, Es, epithelial somite, NT, neural tube, No, notochord. Bar= 50μm.

**Supplementary Fig. S3.**
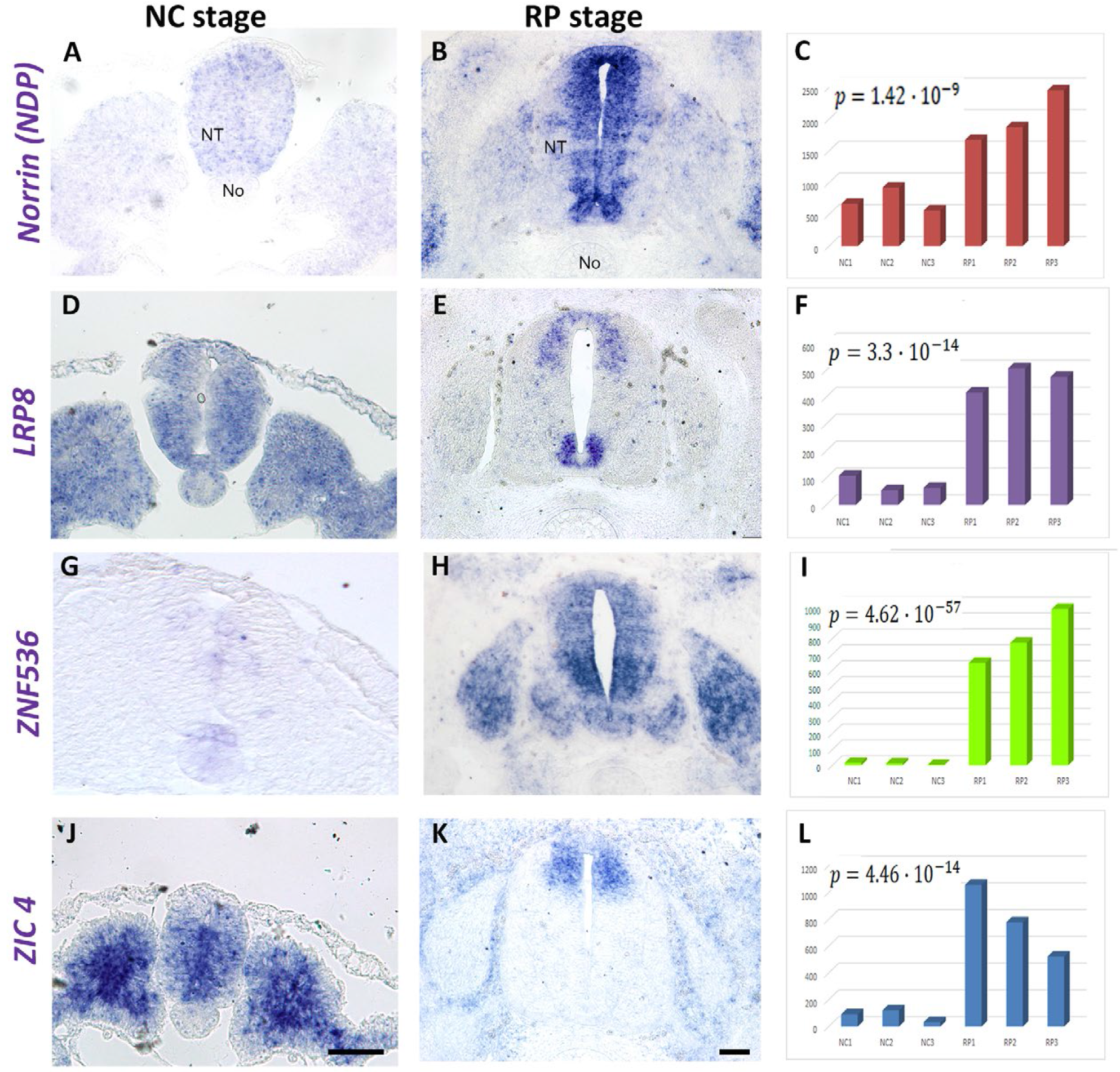
ISH for selected genes expressed in the RP as well as in more ventral domains of the NT. (A, D, G, J) ISH at NC and RP (B, E, H, K) stages, respectively. (C,F,I,L) Quantification of gene expression levels stemming from the transcriptome analysis. In each chart, the B-H adjusted p value is indicated. Note at RP stage the extended expression of transcripts beyond the RP domain. Abbreviations, NT, neural tube, No, notochord. Bar= 50μm.

**Supplementary Fig. S4.**
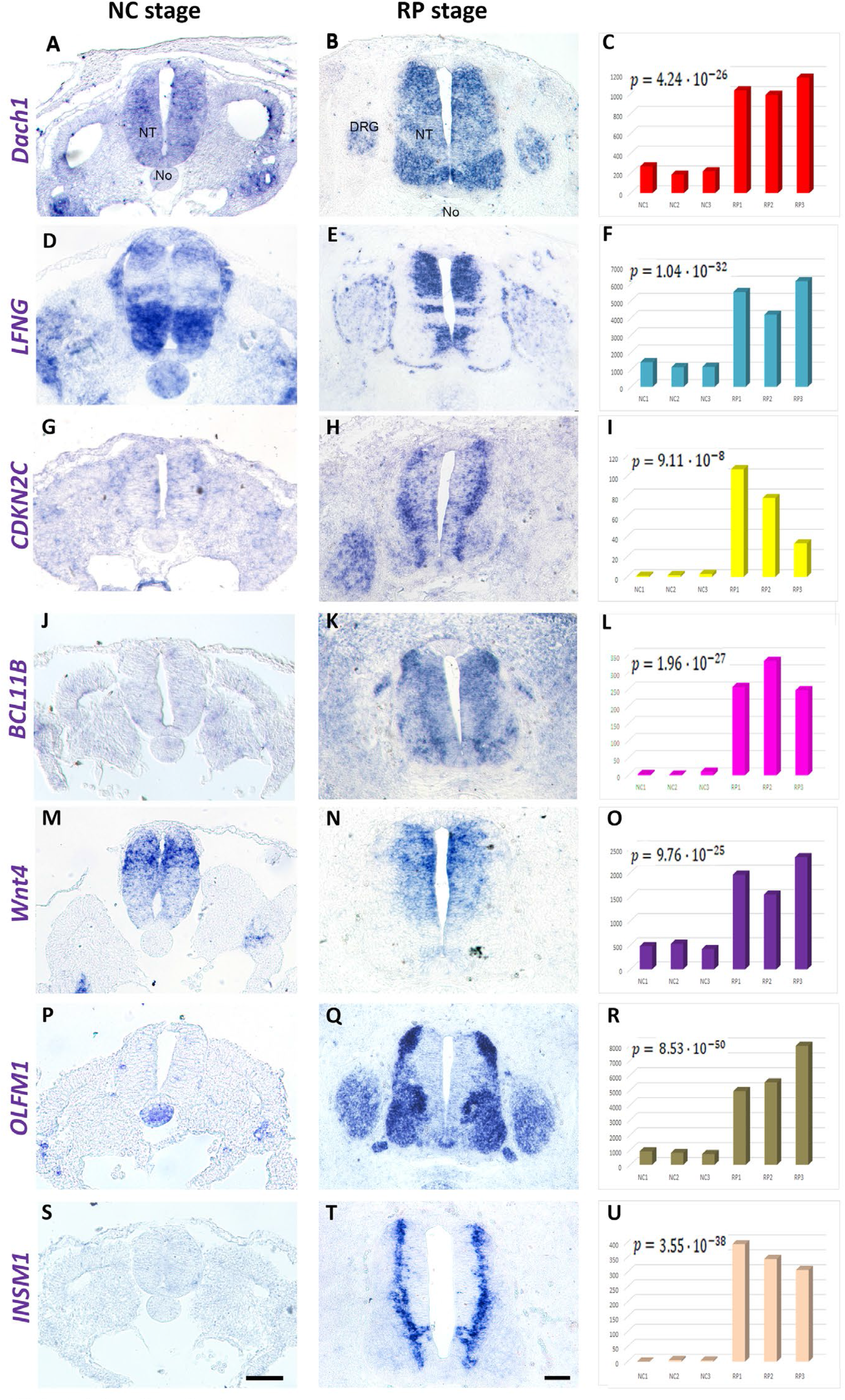
ISH for selected “anti RP” genes upregulated at the RP stage but not transcribed in RP. (A,D,G,J,M,P,S) ISH at the NC stage. (B,E,H,K,N,Q,T) ISH at the RP stage. (C,F,I,L,O,R,U) Quantification of gene expression levels. In each chart, the B-H adjusted p value is indicated. Note in the middle column, the expression of transcripts throughout the NT except for the RP in spite of transcript levels being upregulated in RP compared to NC (right column). Bar= 50 μm.

**Supplementary Fig S5.**
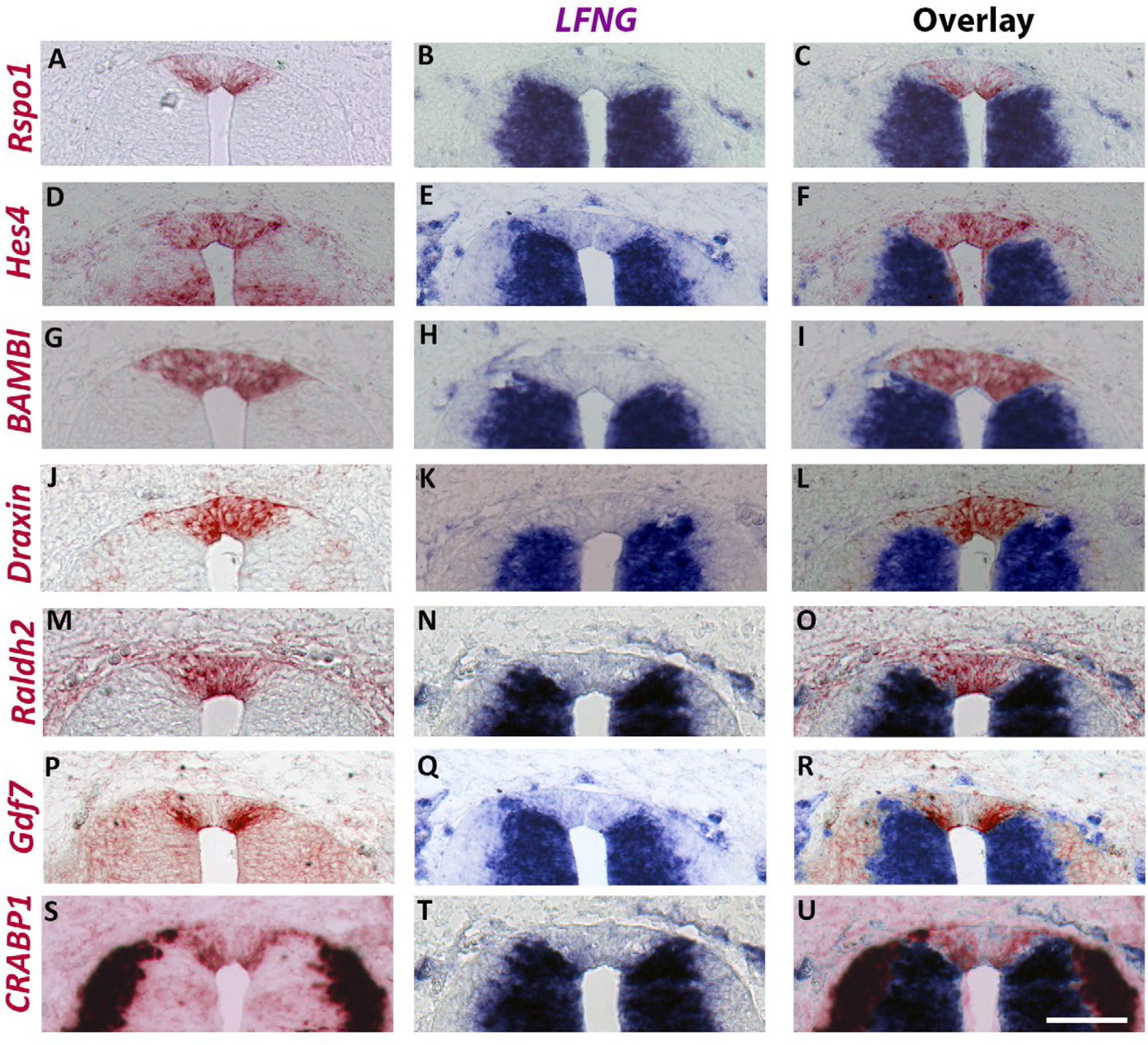
ISH of adjacent sections with selected RP markers in combination with the “anti RP” marker lunatic fringe (LFNG). (A,D,G,J,M,P,S) ISH for RP markers. (B,E,H,K,N,Q,T) ISH for *LNFG*. (C,F,I,L,O,R,U) Overlay of the precedent showing complementary marker expression. Bar= 50μm.

**Supplementary Fig S6.**
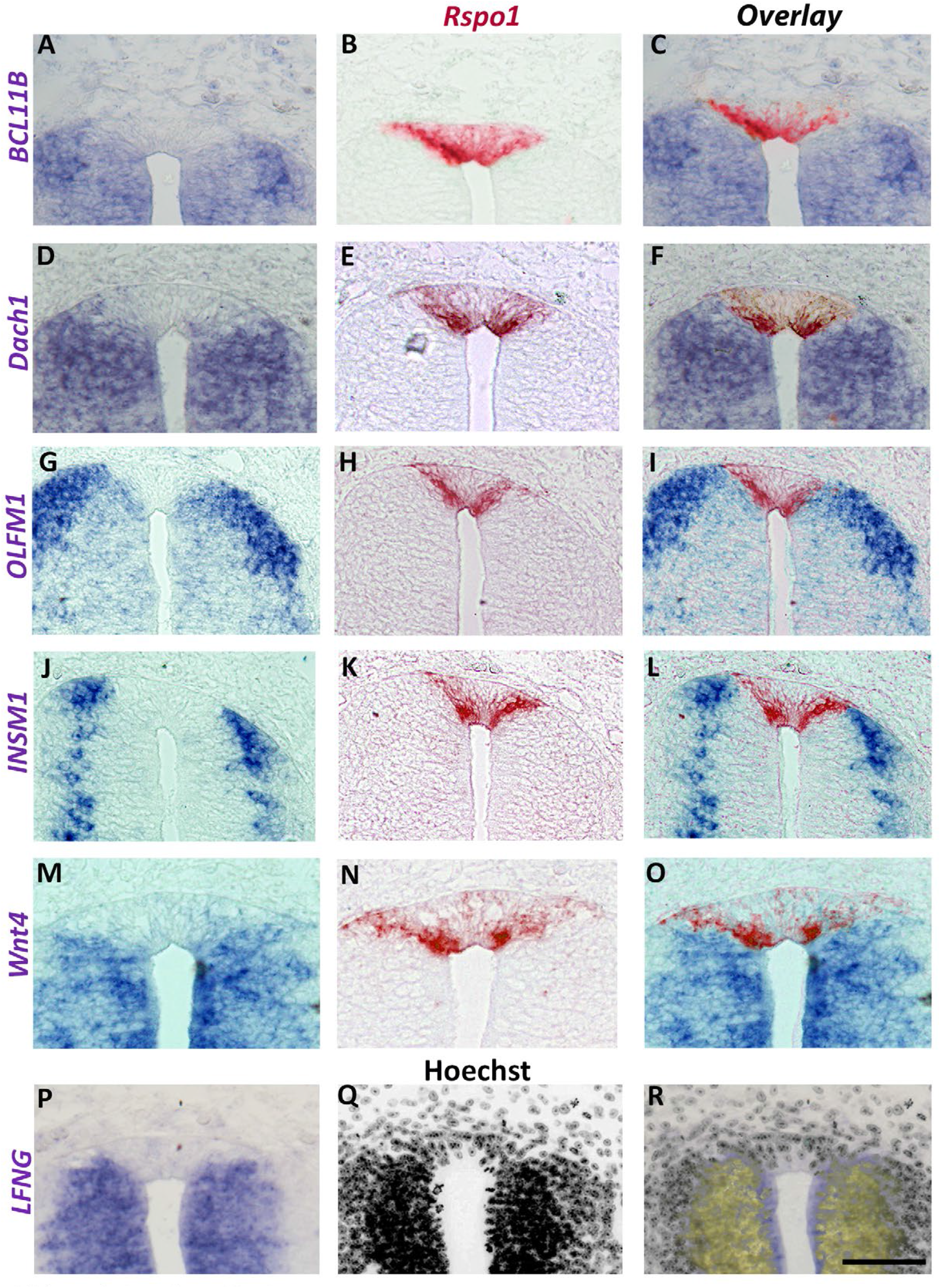
**ISH of adjacent sections with selected “anti RP” genes in combination with Rspo-1**. (A,D,G,J,M) ISH for anti RP markers. (B,E,H,K,N) ISH for *Rspo1*. (C,F,I,L,O) Overlay of the precedent showing complementary marker expression. (P,Q,R) Combined ISH for *LFNG* and Hoechst nuclear staining. Bar= 50μm.

**Supplementary Fig. S7.**
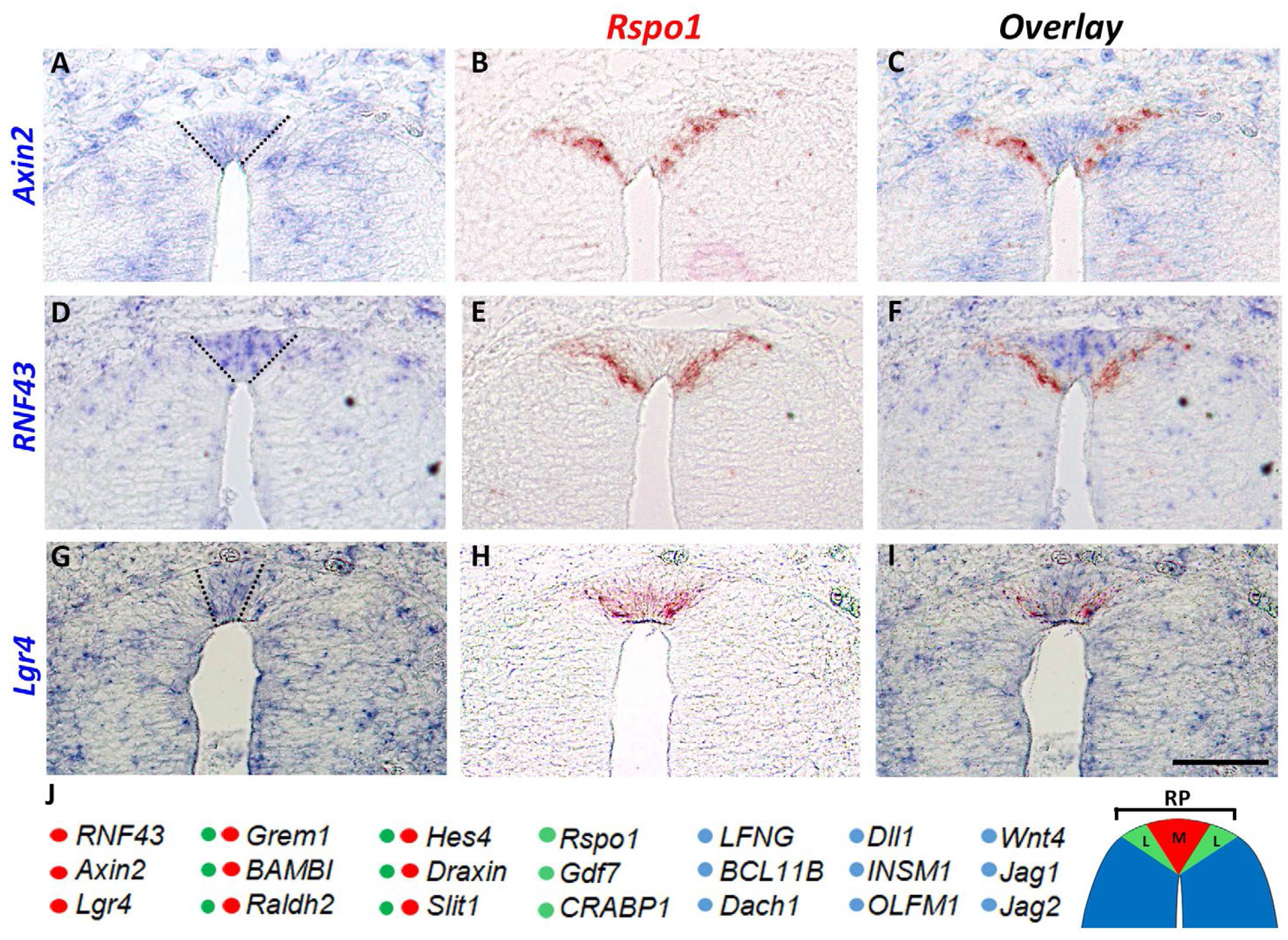
Molecular heterogeneity within the RP. (A,D,G) Expression of three genes in the central domain of the RP (delimited by dotted lines). (B,E,H) *Rspo1* is preferentially expressed in the RP periphery. (C,F,I) Combination of adjacent sections showing complementary expression of the above. (J) Schematic representation of the expression pattern of several genes upregulated at the RP stage to either its medial (M, red), lateral (L, green), to both domains (green+red) or to NT regions except for the RP (blue). Bar= 50μm.

**Supplementary Fig. S8.**
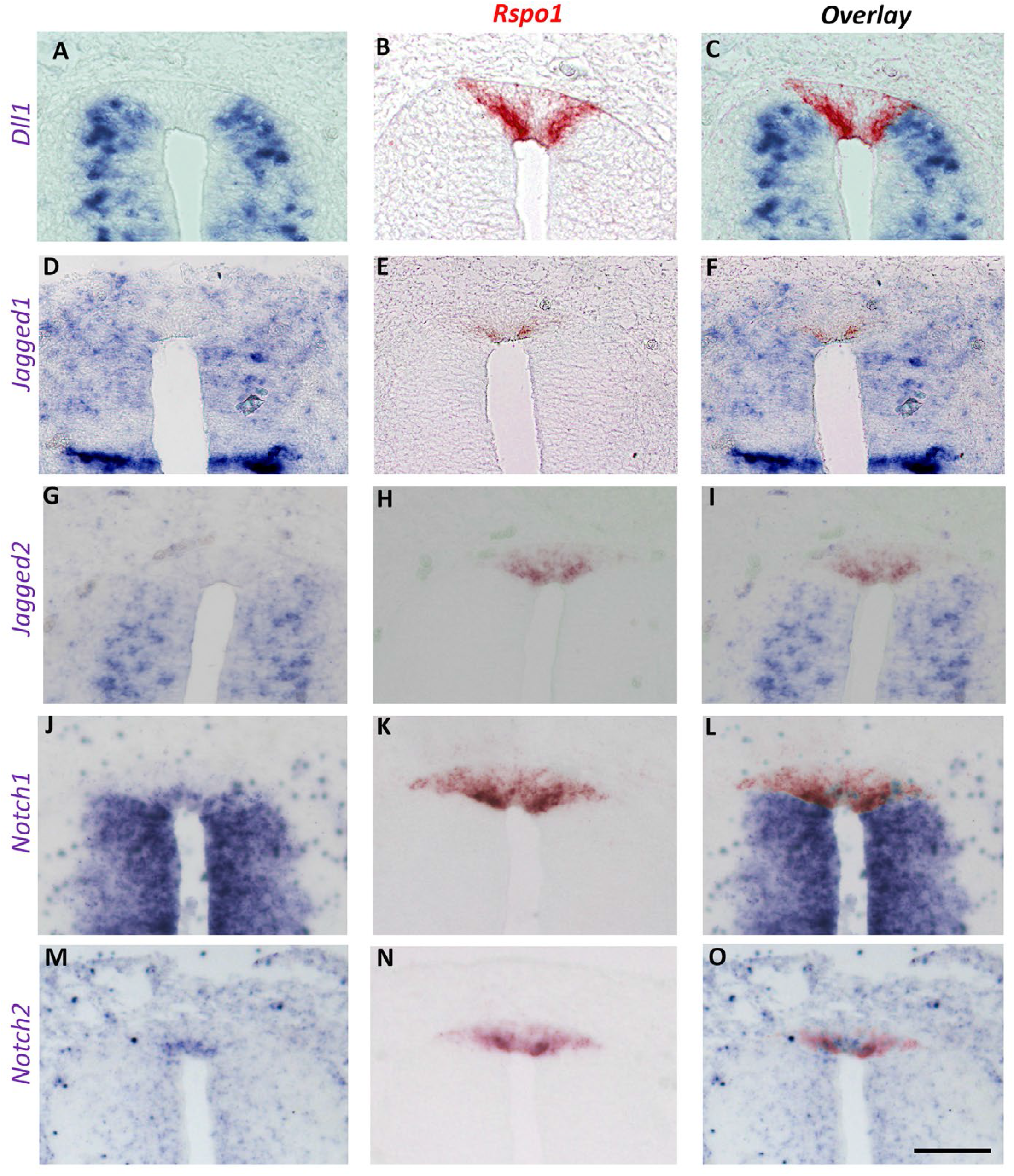
Expression patterns of Notch ligands and receptors. (A,D,G,J,M) Expression of three Notch ligands (A,D,G) and the *Notch1* and *2* receptors (J,M) at the RP stage. (B,E,H,K,N) *Rspo1* expression in adjacent sections. (C,F,I,L,O) Overlay of the above. Note that the dorsal limit of *Dll1*, *Jagged1* and *Jagged2* mRNA expression corresponds to the ventral limit of the RP. *Notch1* and *Notch2* mRNAs are expressed apically in the RP domain. Bar= 50μm.

**Supplementary Fig. S9.**
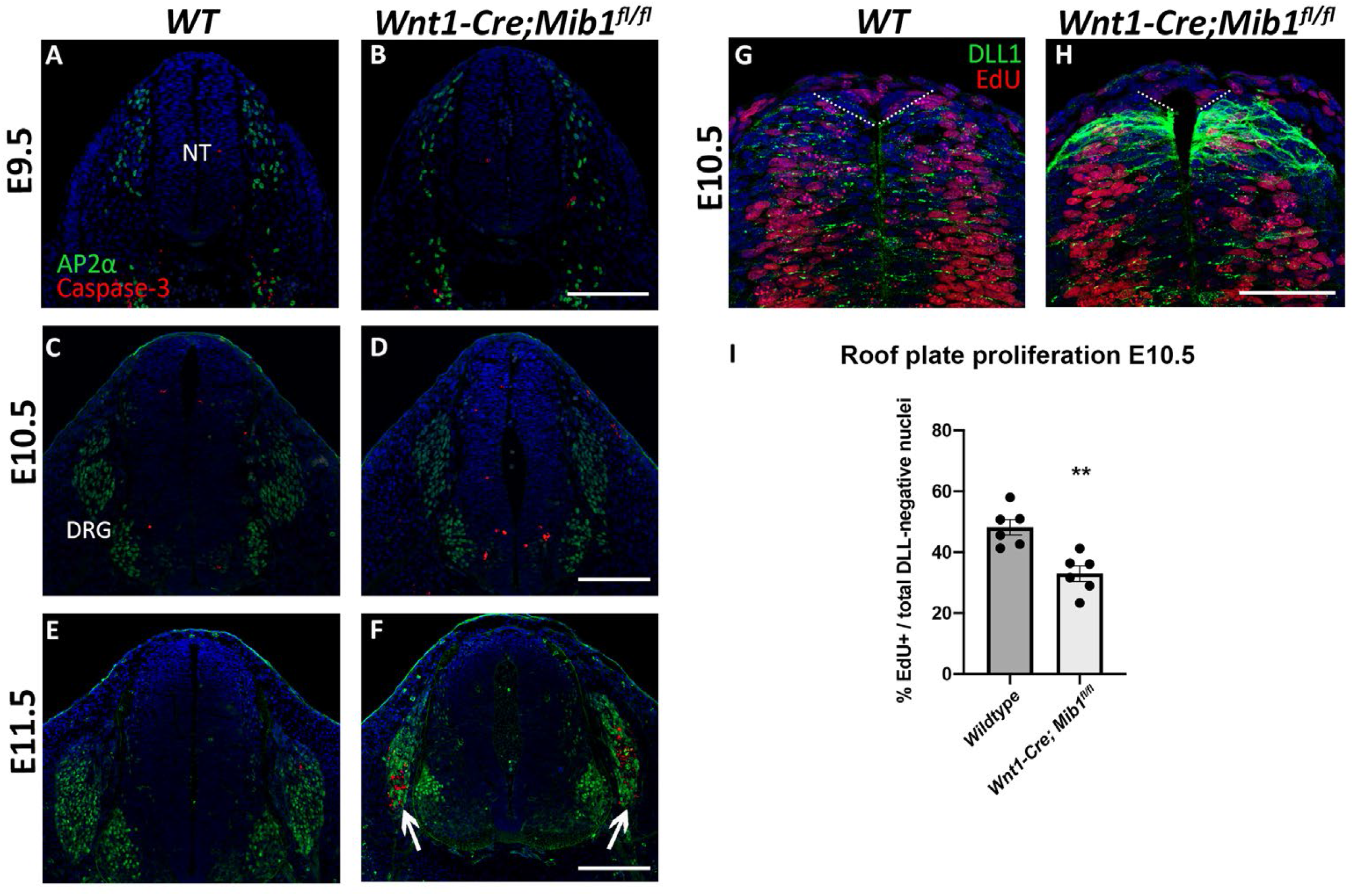
Neural crest formation is preserved in the absence of Notch signaling. (A-F) Transverse sections through the dorsal neural tube at the level of the forelimb in wildtype (WT) and *Wnt1-Cre; Mib1^fl/fl^* mouse embryos, immunostained for AP2*α* labeling neural crest and dorsal root ganglion (DRG) neurons (green), and cleaved-caspase 3 labeling apoptotic cells (red). At all stages examined, no cell death is evident in the premigratory neural crest or presumptive roof plate region. At E11.5 (E, F) increased cell death is evident in the DRG of *Wnt1-Cre; Mib1^fl/fl^* embryos as previously reported (arrows). Bar = 100 μm. (G,H) Transverse sections at E10.5 immunostained for DLL1 (green) and EdU (red) following a 1hr EdU pulse to label proliferating cells. The dashed line indicates the boundary of DLL1 expression and defines the region used for quantification in (I). (I) Quantification of the number of EdU-positive proliferating cells in the Dll1-negative domain (presumptive roof plate) at E10.5, expressed as a percentage of the total number of DAPI-positive nuclei. Note the reduction in proliferating cells present in *Wnt1-Cre; Mib1^fl/fl^* embryos compared to wildtype. N=6 embryos, **p=0.0018. Bar = 50μm.

**Supplementary Table S1.** A list of transcripts downregulated in RP when compared to NC. Excel file containing the list of transcripts downregulated in RP when compared to the premigratory NC stage. Based on triplicate samples for each stage. See Methods for technical details.

**Supplementary Table S2.** A list of transcripts upregulated in RP when compared to NC. Excel file containing the list of transcripts enriched in the RP when compared to the premigratory NC stage. Based on triplicate samples for each stage. See Methods for technical details.

**Supplementary Table S3.** List of primers used for both quail and mouse *in situ* hybridization.

